# Pharmaceutical agent cetylpyridinium chloride modulates mitochondrial dynamics to enhance calcium-stimulated immune mast cell function

**DOI:** 10.64898/2026.11.25.769240

**Authors:** Dylan J. Wagner, Esther Biro, Sydni M. Plummer, Madeleine K. Lorenger, Bright Obeng, Morgan N. Tasker, Jeongwon Eom, Chamithri Weerakoon, Maeve C. Crowley, Emily L. Ledue, Emma R. Morrison, Jonathan F. Sutker, Samuel T. Hess, Julie A. Gosse

**Affiliations:** Department of Molecular & Biomedical Sciences, University of Maine, Orono, ME, USA

**Keywords:** cetylpyridinium chloride, quaternary ammonium compound, mast cell, calcium ionophore, mitochondrial morphology, mitochondrial translocation

## Abstract

The public is routinely exposed to quaternary ammonium compound cetylpyridinium chloride (CPC) through myriad personal care, food, and cleaning products. However, at exposure-relevant and non-cytotoxic doses, CPC exerts mitochondrial- and neurotoxicity, even crossing the blood brain barrier. Previously, our lab showed that CPC inhibits IgE-mediated mast cell function (degranulation) by hampering early tyrosine phosphorylation events and signaling of the lipid PIP_2_, though direct CPC effects on the downstream portion of the pathway remained unknown. We used Ca^2+^ ionophore to directly shuttle Ca^2+^ into RBL-2H3 mast cells, bypassing the upstream events of IgE-mediated signaling and modeling alternative mast cell activation modes. Unexpectedly, CPC causes powerful stimulation of mast cell signaling under inducement of signaling proximal to Ca^2+^ entry; thus, immune pathways alternative to the classic IgE-mediated pathway may be activated by this compound. This finding demonstrates that CPC inhibition of IgE-mediated mast cell signaling solely involves events upstream of Ca^2+^. CPC’s enhancement of Ca^2+^ ionophore-stimulated degranulation is dependent on energy status, perhaps a modification of hormesis. CPC’s mechanism of boosting Ca^2+^ ionophore-mediated degranulation is characterized by effects on Drp1 phosphorylation, mitochondrial fission, energization (increased mitochondrial membrane potential), and mitochondrial translocation, mediated by microtubules. This may be the first instance of a chemical amplifying mitotranslocation. This work links and clarifies connections between Drp1, mitochondrial fission/translocation, and degranulation in basic cell biology. This work expands understanding of CPC’s mitotoxic mechanisms, connecting to its immunomodulatory effects and suggesting impacts of other quaternary ammonium compounds, to ultimately aid consumers and regulators in decision-making.

## Introduction

Consumers are increasingly and regularly exposed (Mohapatra et al. 2023) to the antimicrobial cetylpyridinium chloride (CPC) at millimolar levels (Frantz 2023) in personal care products, cleaning solutions, pharmaceuticals, and food processing (reviewed in Obeng et al. 2024). Quaternary ammonium compounds constitute a broad (and understudied) class of chemicals structurally defined by a positively-charged nitrogen-containing headgroup and hydrophobic tail(s) (Arnold et al. 2023). CPC lyses bacteria at levels above its critical micelle concentration (∼900 µM), at which it acts as a detergent (Mandal and Nair 1991; Varade et al. 2005; Abezgauz et al. 2010; Shi et al. 2011; Mukerjee and Mysels). In our study, all described CPC effects are not due to cellular lysis: doses employed were far lower than the CMC, did not alter plasma membrane integrity, and did not cause cytotoxicity (Raut et al. 2022; Obeng et al. 2023; Weller et al. 2024). The Food and Drug Administration effectively banned CPC specifically from hand soaps and sanitizers (Kux 2016; Gottlieb 2019) but not from many other products. CPC has been exempted from toxicity testing by the U.S. Environmental Protection Agency, but California has elevated CPC as a priority chemical for biomonitoring (Biomonitoring California 2026).

Despite widespread use, we have found no direct data on the human burden of CPC (PubMed, 2026), aside from one small study indicating its presence in human fluids (Environmental Working Group). However, related quaternary ammonium compounds have been detected in human subjects (Zheng et al. 2022), including at levels causing mitochondrial toxicity (Hrubec et al. 2021). To fill this gap, we estimated CPC levels, calculating that daily consumer exposures may reach 1 mg/kg, leading to low- or sub-micromolar levels in human blood, from one-time usage of personal care products and consumption of CPC-treated chicken (Weller et al. 2024). Potential bioaccumulation through chronic use could push levels higher. Evidence for CPC bioaccumulation includes persistence in the body for more than one day following brief exposure of human oral tissues (Bonesvoll and Gjermo 1978) or of mice (Pottel et al. 2020). Following oral product exposure, CPC is slowly released back into saliva over >1 day; this re-released CPC will be ingested, causing ongoing dosing (Bonesvoll and Gjermo 1978). Consumer product exposure-relevant doses of CPC (orally dosed 1 mg/kg) in two mouse studies entered the bloodstream (Pottel et al. 2020) and crossed the blood-brain barrier (Cohn et al. 2024), evincing bioavailability (Voutchkova et al. 2010; Van Leeuwen et al. 2015; Klaassen and Casarett 2019). For example, CPC was found in the mouse brain at nanomolar levels and caused harm to glial cells (Cohn et al. 2024). Also, in mice, CPC rose to blood concentrations within the range of activity against the known target dopamine D3 receptor (Pottel et al. 2020).

Additionally, CPC is toxic to mitochondria (Saladino et al. 1971; Chávez and Bravo 1982; Datta et al. 2017; Liu et al. 2023; Weller et al. 2024; Ledue et al. 2026). CPC potency is similar or higher than various canonical mitotoxicants: carbonyl cyanide 3-chlorophenyl hydrazone, triclosan (Weatherly et al. 2016), 2,4-dinitrophenol (Weller et al. 2024), rotenone, and cyanide (Ledue et al. 2026). CPC causes mitochondria to undergo fission (Weller et al. 2024). A key protein responsible for fission is Drp1 (Zhang et al. 2011; Simula et al. 2018; Kamerkar et al. 2025), though CPC’s mechanism remains unknown.

CPC also impairs the function of mast cells (Obeng et al. 2023; Raps et al. 2024; Weller et al. 2024), present in most tissues including brain and environmental surfaces like the gastrointestinal tract (Kuby 1997). Mast cells are involved in immune and neurological function (Metcalfe et al. 1997; Galli et al. 2005; Silver and Curley 2013), by releasing their specialized granules (degranulation) containing a cocktail of biochemical effectors including histamine and serotonin. The RBL-2H3 line is a widely-used mast cell model which simulates human primary mast cell responses to stimulants and toxicants (Metzger et al. 1982; Seldin et al. 1985; Metcalfe et al. 1997; Abramson and Pecht 2007; Thrasher et al. 2013; Alsaleh et al. 2016).

Mast cells are activated via multiple routes, one being the cell surface high-affinity IgE receptor FcεRI, which becomes crosslinked by incoming foreign antigen. A similar signaling pathway is shared by other immune cells (Marano et al. 1993; Gauld et al. 2002; Smith-Garvin et al. 2009; Trebak and Kinet 2019). Previously, we showed that CPC inhibits mast cell degranulation (Raut et al. 2022; Obeng et al. 2023), though this initial crosslinking event is unaffected (Obeng et al. 2024).

Receptor aggregation results in a tyrosine phosphorylation cascade (Fig. 1) (Kinet 1999) which is inhibited by CPC (Obeng et al. 2024), particularly at Syk kinase and some of its targets like LAT. Activated phospholipase Cγ1 (PLCγ1) cleaves signaling lipid phosphatidylinositol 4,5-bisphosphate (PIP_2_) (Fig. 1), but CPC interferes with PIP_2_ directly (Raut et al. 2022; Aho et al. 2025). Second messengers from PIP_2_ activate store-operated Ca^2+^ entry (SOCE) (Putney 1986; Berridge 1993), flooding the cytosol with sufficient Ca^2+^ to activate enzymes and microtubules. Newly-polymerized microtubules deliver granules to the plasma membrane for exocytosis (Fig. 1) (Nishida et al. 2005). SOCE and degranulation are inhibited by CPC (Obeng et al. 2023).

**Figure 1.**
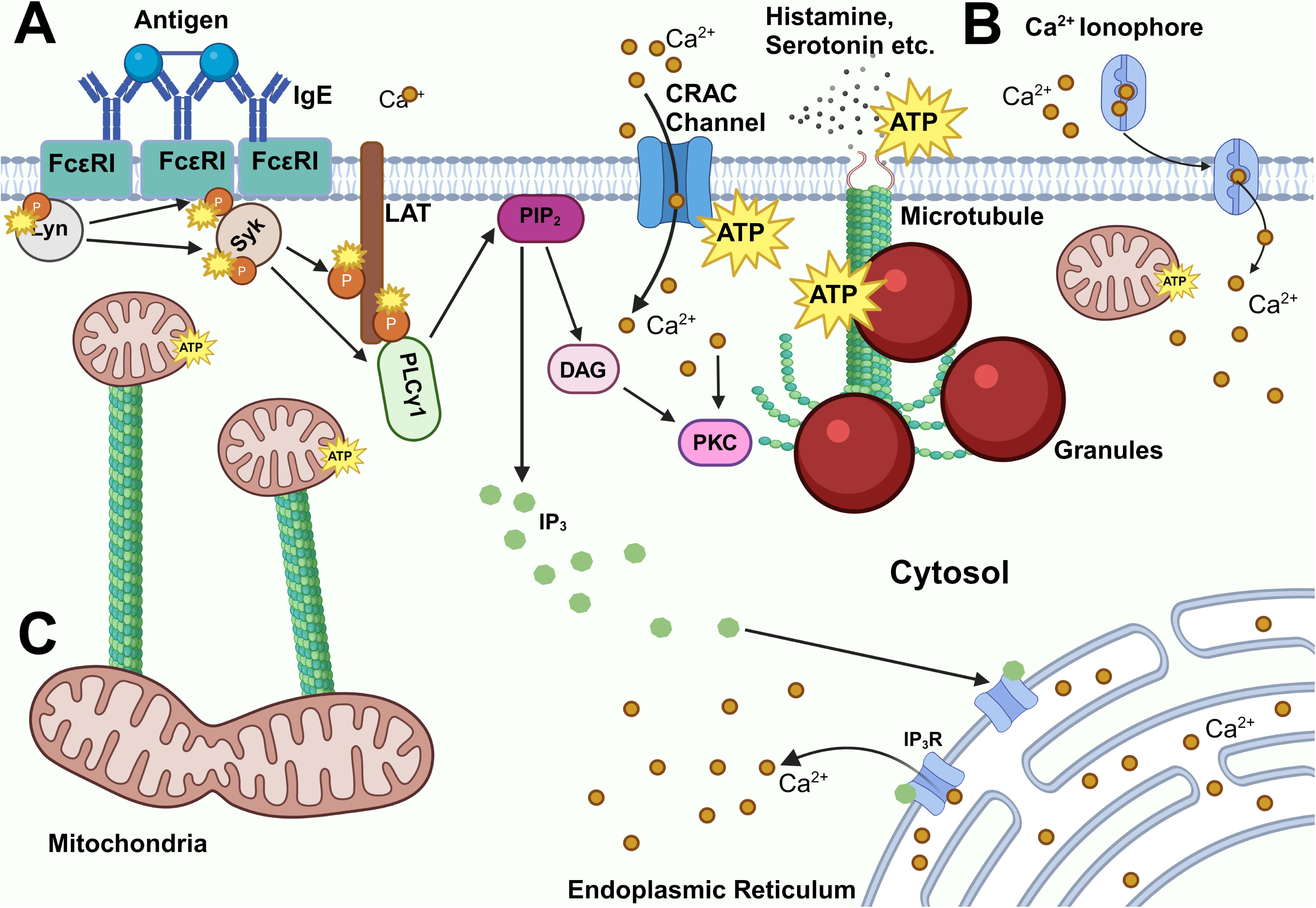
Antigen-mediated and Ca^2+^ ionophore-mediated signaling pathways in mast cells. **(A)** Foreign antigen binds to IgE-bound FcεRI receptors, crosslinking them. Crosslinking of the FcεRI receptors leads to an early tyrosine phosphorylation cascade involving signaling proteins Lyn kinase, Syk kinase, adaptor LAT, and phospholipase PLCγ. Small yellow bursts indicate an activating phosphorylation event. Activated PLCγ cleaves PIP_2_ into secondary messengers diacylglycerol (DAG) and inositol 1,4,5-triphosphate (IP_3_). Subsequently, IP_3_ (green circles) binds to the IP_3_ receptor on the endoplasmic reticulum membrane, releasing internal Ca^2+^ (brown dots) stores, thus triggering SOCE into the cell. SOCE is mediated by the endoplasmic reticulum in collaboration with cell surface Ca^2+^ release-activated Ca^2+^ (CRAC) channels. Large yellow bursts indicate use of ATP. Increasing cytosolic Ca^2+^ levels trigger microtubule polymerization organized by protein kinase C (PKC) and, finally, the release of internal granules. These granules (large red bubbles) fuse to the plasma membrane, exocytosing their biochemical effector contents, such as histamine and serotonin. **(B)** Calcium ionophore shuttles Ca^2+^ directly into the cell, completely bypassing all events prior to SOCE but still triggering microtubule polymerization and degranulation. **(C)** Mitochondria (small yellow bursts indicate ATP production) move from the perinuclear space towards the cellular periphery in order to aid the energy-intensive process of degranulation.

Drp1-driven mitochondrial fission leads to mitochondrial translocation from the perinuclear space towards the plasma membrane in mast cells (Fig. 1) (Zhang et al. 2011); a similar mitochondrial translocation occurs in activated T cells (Quintana et al. 2007).

While CPC is known to directly inhibit Syk phosphorylation (Obeng et al. 2024) and PIP_2_ (Raut et al. 2022; Aho et al. 2025), it is unknown whether CPC targets components farther downstream than PIP_2_. To understand the full suite of CPC targets in immune signaling, we utilized an experimental tool, Ca^2+^ ionophore, to directly shuttle Ca^2+^ into the cell (Fig 1) (Foreman et al. 1973; Kagayama and Douglas 1974; Shim, Weatherly, et al. 2016). Thus, we set out both to understand CPC’s effects on signaling post-SOCE and to simulate non-IgE pathways of mast cell activation, which are also physiologically important. We discovered an interesting phenomenon caused by this widespread antimicrobial – stimulated mast cell function – and then investigated the underlying mechanism, focusing on the interplay between mitochondria and immune signaling.

## Materials and Methods

Safety precautions, per each chemical’s safety data sheet, were used for each chemical and assay.

### Chemical Preparation

Cetylpyridinium chloride monohydrate (CPC; 99.7% purity; MP Biomedicals, CAS no. 6004-24-6) was prepared by dissolving into Tyrodes buffer, pH 7.4, with 1 mg/mL bovine serum albumin (BSA; low heavy metals, 98% purity, Millipore Sigma) (BT) (Hutchinson et al. 2011). CPC was dissolved without the use of organic solvents, and concentration was verified via spectrophotometry (Bernauer et al. 2015; Raut et al. 2022).

Calcium ionophore (Millipore Sigma, CAS no. 52665-69-7) was reconstituted in DMSO. Thus, “Spont.” samples in ionophore experiments received 0.0032% DMSO in the vehicle control.

### RBL-2H3 Mast Cell Culture

RBL-2H3 mast cells were cultured as described (Hutchinson et al. 2011). Cells for microscopy experiments were cultured in phenol red-free media (with no L-glutamine; Gibco), with added fetal bovine serum (GeminiBio), gentamicin (Gibco), and L-glutamine (Gibco). Mycoplasma tests were performed to ensure cells were free of contamination.

### Degranulation Assay

RBL-2H3 mast cell degranulation was measured using the assay described (Obeng et al. 2023). All CPC treatments were conducted in BT for 1 hr, concurrent with ionophore exposure. Ionophore dose was chosen as 300 nM, to balance the need for sufficiently robust signaling (Fig. S1) over spontaneous levels with the goal of avoiding maximally-stimulating or cytotoxic (Fig. S2) doses.

### Western Blotting

Proteins were probed using the western blot protocol described in (Obeng et al. 2024) with the following modifications. Drp1 and phospho-Drp1 Ser616 (Cell Signaling Technologies) antibodies were used, each at 1:1,000 dilution in 1X TBST with 5% BSA. Cells were treated with CPC for 1 hr, but no ionophore or antigen stimulation was used. The amount of protein used to load into each lane was within the linear response range for each antibody (Fig. S5). Using our published methodology (Obeng et al. 2024), blots of identical samples run on and transferred from the two sides of the gel were cut in two for direct comparison of phospho- to total protein. The secondary antibody utilized was IRDye 680RD Goat Anti-Rabbit IgG (LI-COR) (1:20,000, diluted in 1% non-fat dry milk-TBS) by incubating at RT for 3 hr, protected from light. Following LI-COR imaging, protein bands of Ser616-phosphorylated Drp1 were normalized to total Drp1 in each sample.

### Confocal Imaging of Mitochondrial Translocation

RBL-2H3 mast cells were plated and incubated as in the mitotranslocation experiments described previously (Weatherly et al. 2018). The following day, cells were incubated with 20 nM MitoTracker Red FM (Thermo Fisher; dissolved in 100% DMSO, resulting in addition of 0.002% DMSO vehicle upon dilution) and 100 nM Hoechst 33342 (stock dissolved in water; Biotium) dyes in BT for 30 min. Thus, ionophore-treatment samples received a total organic solvent vehicle of 0.0052% DMSO, a level previously shown to not affect mitochondrial and immune signaling function in RBL-2H3 cells (Weatherly et al. 2016).

If cells were to receive antigen treatment (methods in Obeng et al. 2023), cells were sensitized with 0.1 µg/mL anti-dinitrophenyl IgE in BT (1 hr) before dye treatment. Then, cells were exposed to either 0.0007 µg/mL dinitrophenyl-BSA antigen (which produced a moderate 20 - 30% degranulation response, data not shown) or 300 nM Ca^2+^ ionophore (see degranulation response in Fig. S1) to activate mast cell degranulation. Concurrently with antigen stimulation, cells received 10 µM CPC, a known inhibitory dose (Obeng et al. 2023) for the 45 min exposure; control cells for antigen experiments received a BT vehicle control. Concurrently with ionophore stimulation, cells received 2.5 µM CPC, the stimulatory dose (Fig. 2), for a 1 hr exposure; control cells for these ionophore experiments received 0.0082% DMSO in total.

**Figure 2.**
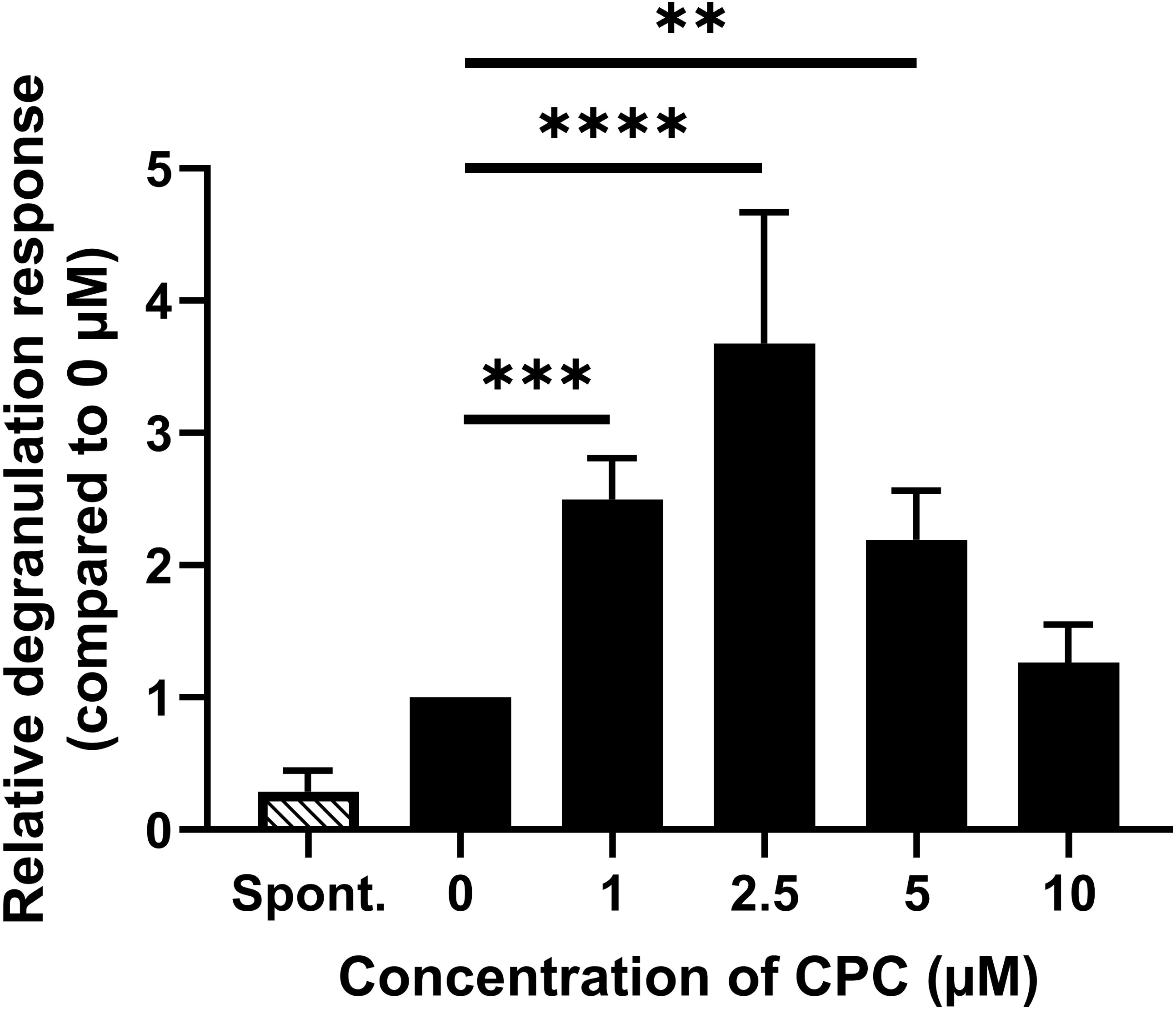
Relative degranulation response of RBL-2H3 mast cells exposed to various doses of CPC with stimulation by Ca^2+^ ionophore. Spontaneous release cells (“Spont.”) were exposed to BT in the absence of Ca^2+^ ionophore or CPC. Sample wells were exposed to 0 to 10 µM CPC along with 300 nM Ca^2+^ ionophore for 1 hr. Values shown are means that were normalized to the control (0 μM CPC/300 nM ionophore), are displayed ± SEM, and represent at nine independent days of experiments (each sample in triplicate). Statistical significance was determined through one-way ANOVA with Tukey’s post-hoc testing: \**p* < 0.1, \*\**p* < 0.05, \*\*\**p* < 0.01, \*\*\*\**p* < 0.001, \*\*\*\*\**p* < 0.0001.

Following these treatments, live cells were imaged on the Zeiss LSM 980 confocal microscope with the Airyscan 2 detector over the course of ∼15 min. Hoechst 3342 fluorescence (Excitation 405 nm, Emission 422 - 477 nm) and MitoTracker Red FM fluorescence (Excitation 561 nm, Emission 570 - 630 nm) were measured with an objective of numerical aperture 1.4, along with differential interference contrast images.

Control experiments were performed to test whether CPC interferes with MitoTracker Red FM (Fig. S7): CPC does not interfere with the fluorescence of MitoTracker Red FM.

### Mitochondrial Translocation Analysis

To determine the percentage of mitochondrial translocation towards the plasma membrane, the area of mitochondria (number of pixels with MitoTracker Red FM signal above background), within the perinuclear region *versus* in the whole cell, were measured. An ImageJ Fiji macro was created for this analysis, and both the code for it and a visual description of the image analysis workflow are available in Fig. S8.

### Analysis of Mitochondrial Morphology

Mitochondrial morphology/networking data were extracted from the same images captured by the methods described above in “Confocal Imaging of Mitochondrial Translocation.” All collected images were analyzed. The Mitochondria Analyzer Fiji Plugin was used to analyze the morphology of mitochondrial elements (∼100-150 per cell) as described (Chaudhry et al. 2020). The 2D Batch Command was used for high-throughput processing, providing the mean value of mitochondrial element features (*e.g.* perimeter) for all of the mitochondrial elements in every image. The recommended pre- and post-processing steps were followed (Chaudhry et al. 2020) with the settings recommended by the authors. A Weighted Mean thresholding method was utilized, starting with a standard Block Size and C-value that were then optimized to both reduce background noise and preserve structural features. Displayed data were for parameters that report measures of mitochondrial networking and shape.

### Confocal Imaging of Microtubule Polymerization

Microtubule polymerization was assessed as previously described (Weatherly et al. 2018; Obeng et al. 2023), using the pIRESneo-EGFP-alpha Tubulin construct (a gift from Patricia Wadsworth; Addgene plasmid #12298; https://www.addgene.org/12298/) (Rusan et al. 2001). Cells were electroporated with this plasmid in phenol red-free RBL-2H3 media using an ECM 630 electroporator (Harvard Bioscience). Electroporation of samples were pulsed once in 2 mm gap cuvettes with the following conditions: 250 V, 950 µF capacitance, 25 Ω resistance, and a desired field strength of 625 V/cm.

Electroporated cells (240,000/well) were plated in 8-well ibidi plates in 200 µL/well of phenol red-free media, and incubated overnight at 37 °C with 5% CO_2_. The next day, cells were washed with BT, then incubated for 30 minutes with 20 nM MitoTracker Red (Thermo Fisher) in BT (200 µL/well). After incubation, cells were washed with BT and dosed with 200 µL/well of either 0.0032% BT-DMSO (control), 300 nM Ca^2+^ ionophore, or 2.5 µM CPC and 300 nM Ca^2+^ ionophore, then incubated for 1 hr at 37°C. Following this incubation, EGFP fluorescence (Excitation 488, Emission 490 - 550 nm) and MitoTracker Red fluorescence (Excitation 561 nm, Emission 570 - 630 nm) in live cells were imaged using the Zeiss Airyscan (see “Confocal Imaging of Mitochondrial Translocation”) for up to 30 min.

### Mitochondrial Membrane Potential

Mitochondrial Membrane Potential (MMP) data were extracted from the same images captured by the methods described in “Confocal Imaging of Mitochondrial Translocation.” All collected images were analyzed. Higher MitoTracker Red FM accumulation in the mitochondrial matrix increases the dye’s fluorescence intensity in the cell, reflecting higher MMP (Buravkov et al. 2014; Weatherly et al. 2018). The mitochondrial region of interest was outlined, and the mean gray value, or average MitoTracker Red FM fluorescence intensity per pixel, was measured. Normalization of signal per pixel prevents changes in cell area (*e.g.*, observed in degranulating cells) from affecting the MMP value. Background signal (taken from a cell-sized region of interest in the image background) was subtracted from each recorded cellular mean gray value.

### Imaging and Statistical Analyses

Numerical data reported in this study are expressed as mean ± SEM. Microscope images were analyzed using Fiji ImageJ2 (v. 2.16.0/1.54). Microsoft Excel (Microsoft 365) and GraphPad Prism (v. 11.1.0) were used to create figures along with performing statistical testing. Statistical analyses were performed with a starting alpha of 0.1 in order to balance Type I and II errors, with the goal to not miss subtle effects in light of the modest ionophore stimulation conditions used in this study.

## Results

### CPC enhances Ca^2+^ ionophore-stimulated degranulation of mast cells

Previously, we discovered that CPC inhibits human and rat mast cell degranulation when the cells are stimulated via IgE receptor activation. In the current study, in order to pinpoint the molecular mechanism of this inhibition, we used the tool of Ca^2+^ ionophore stimulation to dissect the pathway into two sections: upstream and downstream of Ca^2+^ influx across the plasma membrane. The method probes whether bypassing early signaling steps known to be inhibited by CPC (tyrosine phosphorylation (Obeng et al. 2024) and PIP_2_ (Raut et al. 2022; Aho et al. 2025) reveals additional inhibitory targets. In order to do so, we first assayed the degranulation dose-response of Ca^2+^ ionophore in glucose-containing BT buffer, and based on these results, we chose an appropriate ionophore dose (300 nM) going forward (Fig. S1). Without CPC present, Ca^2+^ ionophore (300 nM) elicits average absolute degranulation responses, respectively, of 7 ± 1% (SEM) (values were 5.1 - 10.8%) degranulation for the control (0 µM CPC with 300 nM ionophore). Spontaneous samples ((“Spont.”; those treated with BT alone, no simulant) evince an absolute degranulation level of 1.9 ± 0.3%. These Ca^2+^ ionophore stimulation conditions are non-cytotoxic, including in conjunction with a range of CPC doses used in this study (Fig. S2).

We then exposed RBL-2H3 mast cells to Ca^2+^ ionophore plus various concentrations of CPC. Unexpectedly, CPC stimulates degranulation. In an inverted U-shaped dose response curve (Fig. 2), degranulation peaks at 2.5 µM CPC, which causes a ∼4-fold increase over ionophore alone. (Note, the same stimulatory phenomenon was observed with 400 nM ionophore, Fig. S9B). However, the effect dissipates by 10 µM CPC.

This synergistic CPC effect on mast cell degranulation was recapitulated in degranulation experiments under other physiologically relevant conditions (Figs. S3, S4). Fig. S3 shows the same stimulatory phenomenon for cells in BT containing galactose (which is typically shunted by the cell for biosynthesis and purposes other than catabolism), in substitution for glucose. Under these conditions, the cells are relatively low on energy but still show CPC-enhanced degranulation, albeit only ∼2-fold (compared to the 4-fold effect in the presence of glucose) (Fig. S3). When the experimental media was changed from buffered saline solution to a glucose-free, galactose-containing DMEM media, the CPC effect on ionophore-stimulated degranulation became more nuanced (Fig. S4). In this situation without any glucose present, the CPC effect varied in a manner dependent on the extent of mast cell degranulation: CPC caused no effect at very low levels of degranulation, some stimulation with low levels of degranulation, and inhibition at moderate levels of granule release. Fig. S4A contains results of experiments in which ionophore-induced stimulation was very low (3.5%, a value only slightly above spontaneous degranulation), a situation comparable to the results found with unstimulated cells in our previously-published work (Obeng et al. 2023). Fig. S4B contains results from experiments in which ionophore-induced stimulation was higher (10.8%, comparable to the level due to ionophore alone in Fig. 2): this situation evinced a hint of the stimulation ∼31% at 2.5 µM) in Fig. 2 but did not reach statistical significance. In contrast, in cells undergoing higher levels of degranulation (average 20.4%) in this galactose DMEM (Fig. S4C), very low doses of CPC actually inhibited mast cell degranulation.

These results are consistent with CPC primarily targeting early tyrosine phosphorylation, PIP_2_, and mitochondrial function (Fig. 1) rather than directly inhibiting events after Ca^2+^ mobilization (Fig. 1). However, the data raise a new question: how does CPC boost mast cell degranulation under Ca^2+^ ionophore stimulation?

### CPC increases phosphorylation at an activating residue within mitochondrial fission protein Drp1

In mast cells, activated Drp1 is required for the fission of mitochondrial networks and subsequent mitochondrial translocation (“mitotranslocation”) in mast cells towards the plasma membrane (Zhang et al. 2011). Our previous experiments indicated that CPC induces mitochondrial fission (Weller et al. 2024). Thus, we hypothesized that CPC-induced mitochondrial fission would lead to increased mitotranslocation, thereby increasing mast cell degranulation as well. To test this hypothesis, we first assessed CPC effects on activation of Drp1, the primary driver of the mitochondrial fission and translocation process (Zhang et al. 2011). Drp1 phosphorylation at Ser616 is necessary for its activation (Taguchi et al. 2007). Accordingly, we investigated whether CPC affects the phosphorylation of this key residue in Drp1 with western blotting (Fig. 3), normalizing results to total Drp1 signal assessed from the same samples. RBL-2H3 cells were exposed for 1 hr to CPC in glucose-containing BT.

**Figure 3.**
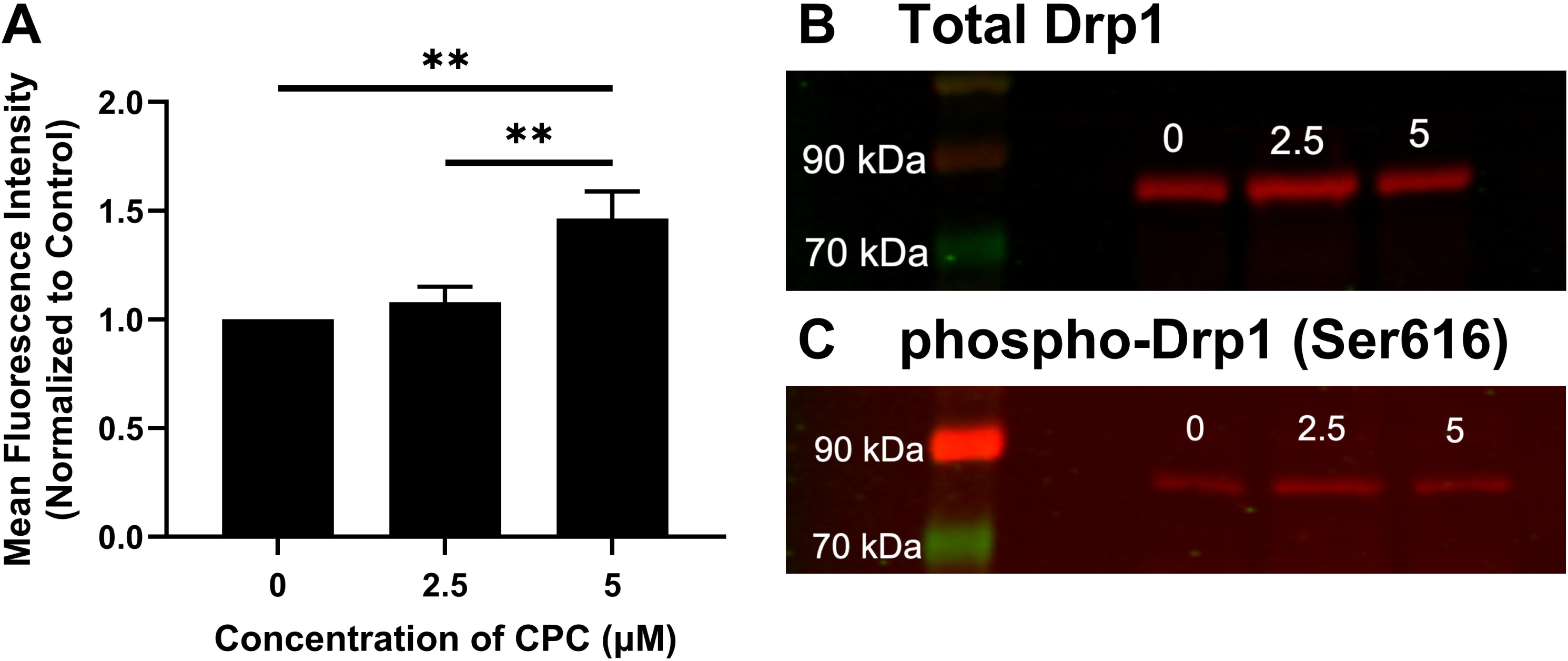
Drp1 phosphorylation (Ser616), assessed via western blot, in unstimulated RBL-2H3 mast cells treated with and without CPC. Cells were treated with 0 µM, 2.5 µM, or 5 µM of CPC for 1 hr before lysis, then probed with total Drp1 or phospho-Drp1 (Ser616) antibodies. On each day, the phospho-Drp1 signal was normalized to the corresponding total Drp1 signal. (**A**) Data are mean ± SEM and represent three independent experiments normalized to the control group (0 µM). Statistical significance was determined through one-way ANOVA with Tukey’s post-hoc testing: \**p* < 0.1, \*\**p* < 0.05, \*\*\**p* < 0.01, \*\*\*\**p* < 0.001, \*\*\*\*\**p* < 0.0001. Total Drp1 (**B**) and phospho-Drp1 (Ser616) (**C**) are representative images of blots. Ladder represents molecular weight in kDa.

Amplification of Drp1 Ser616 phosphorylation is observed due to CPC exposure, reaching statistical significance at 5 µM (1.5-fold increase) (Fig. 3). These data demonstrate that CPC activates the serine phosphorylation at residue 616 of Drp1.

In order to investigate another potential Drp1 regulatory site, we assessed phosphorylation at Ser637. We found that phosphorylation at Drp1 Ser637 is very low in RBL-2H3 mast cells, whether resting or treated with CPC (Fig. S6).

### CPC stimulates mitochondrial translocation in Ca^2+^ ionophore-stimulated mast cells

To test the hypothesis that CPC enhances the translocation of mitochondria to the plasma membrane in Ca^2+^ ionophore-stimulated mast cells, we utilized MitoTracker Red FM to label mitochondria and Hoechst 33342 to label nuclei of RBL-2H3 mast cells in glucose-containing BT (Fig. 4). We found that in resting mast cells (“Spont.”), most of the mitochondria in the cell reside in the perinuclear space (Figs. 4A, D), with only 33 ± 1% (mean ± SEM) residing outside of this space. Addition of 300 nM Ca^2+^ ionophore for 1 hr, as utilized in the degranulation experiments, increases the level of translocated mitochondria (away from the perinuclear space) to 44 ± 2% (Figs. 4B, D). Treating with both Ca^2+^ ionophore and 2.5 µM CPC (the peak dose for enhancing degranulation, Fig. 2) further increases translocated mitochondria to 51 ± 2% (Figs. 4C, D). These data led to the hypothesis that CPC increases mitochondrial translocation in ionophore-stimulated cells, as a mechanism underlying the increased degranulation.

**Figure 4.**
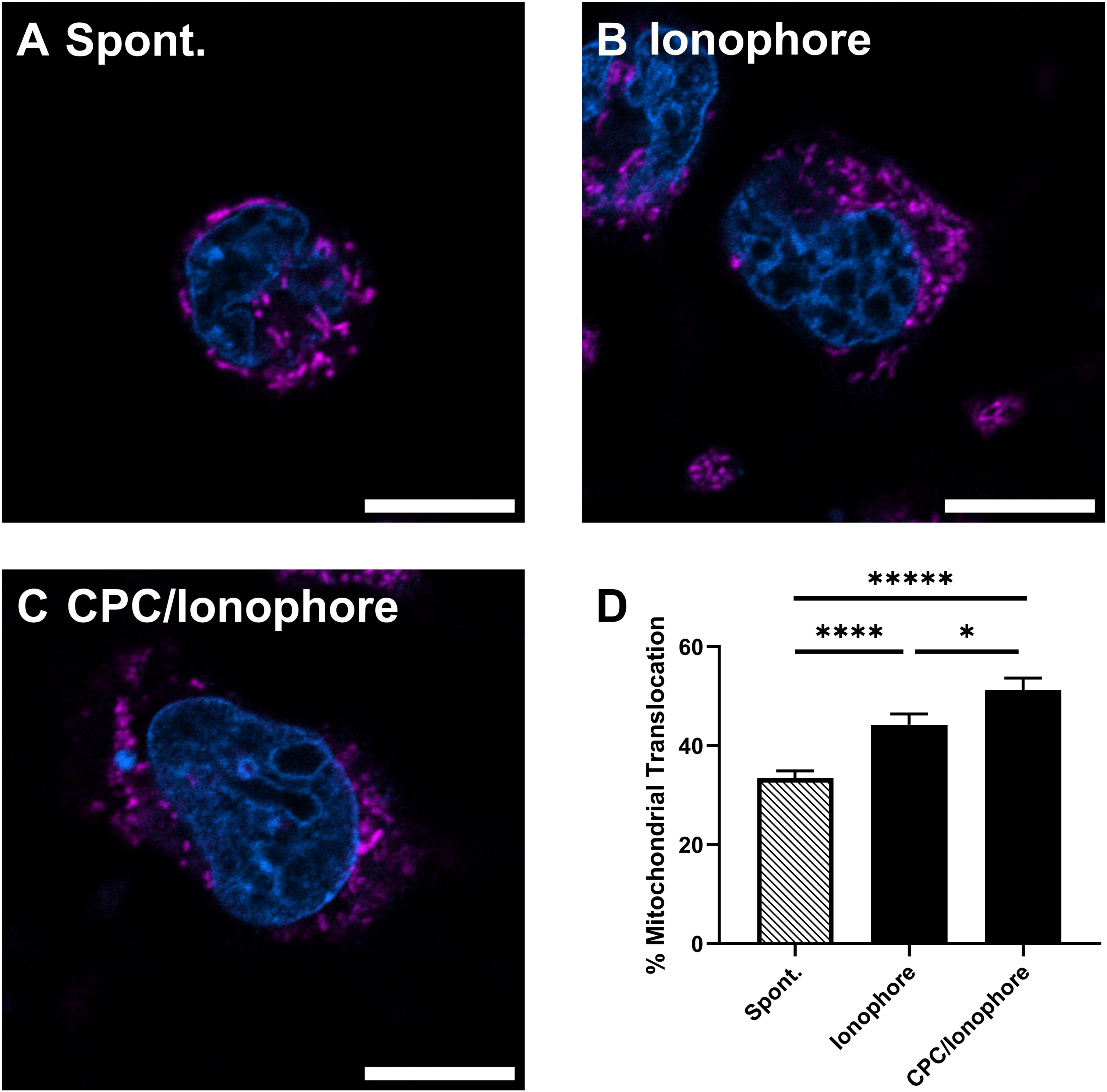
Mitochondrial translocation in ionophore-stimulated RBL-2H3 mast cells treated with or without CPC. To assess mitochondrial position, cells were first stained with Hoechst 3342 (nucleus; blue) and Mitotracker FM (mitochondria; magenta) as described in Materials and Methods, then washed and placed into BT with the stated treatments for 1 hr, followed by immediate Airyscan imaging of live cells within ∼15 min. Representative images of cells treated with **(A)** only BT (“Spont.”; N = 179 cells), **(B)** 0 µM CPC + 300 nM ionophore (N = 69), or **(C)** 2.5 µM CPC + 300 nM ionophore (N = 55). Scale bar represents 10 µm. **(D)** Quantification of mitochondrial localization away from the nucleus and toward the plasma membrane. Data are mean ± SEM and represent four independent experiments. Statistical significance was determined through one-way ANOVA with Tukey’s post-hoc testing: \**p* < 0.1, \*\*\*\**p* < 0.001, \*\*\*\*\**p* < 0.0001.

### CPC inhibits mitochondrial translocation in antigen-stimulated mast cells

In contrast to Ca^2+^ ionophore stimulation conditions, CPC inhibits (does not enhance) degranulation during antigen stimulation of IgE receptors (Obeng et al. 2023); thus, we also measured mitotranslocation under these conditions, to put the ionophore findings into context. We utilized a mast cell inhibitory dose of CPC (10 µM) (Obeng et al. 2023). We found that in resting mast cells (“Spont.”), most of the mitochondria in the cell reside in the perinuclear space (Figs. 5A, D), with only 30 ± 2% residing outside of this space, similar to Fig. 4A. Addition of 0.0007 µg/mL antigen increased the level of translocated mitochondria (away from the perinuclear space) to 47 ± 2% (Figs. 5B, D), a similar extent of translocation as did ionophore (Fig. 4B, D). Addition of CPC (with antigen stimulation) inhibits the mitotranslocation, back down to control levels: 35 ± 2% (Figs. 5C, D).

**Figure 5.**
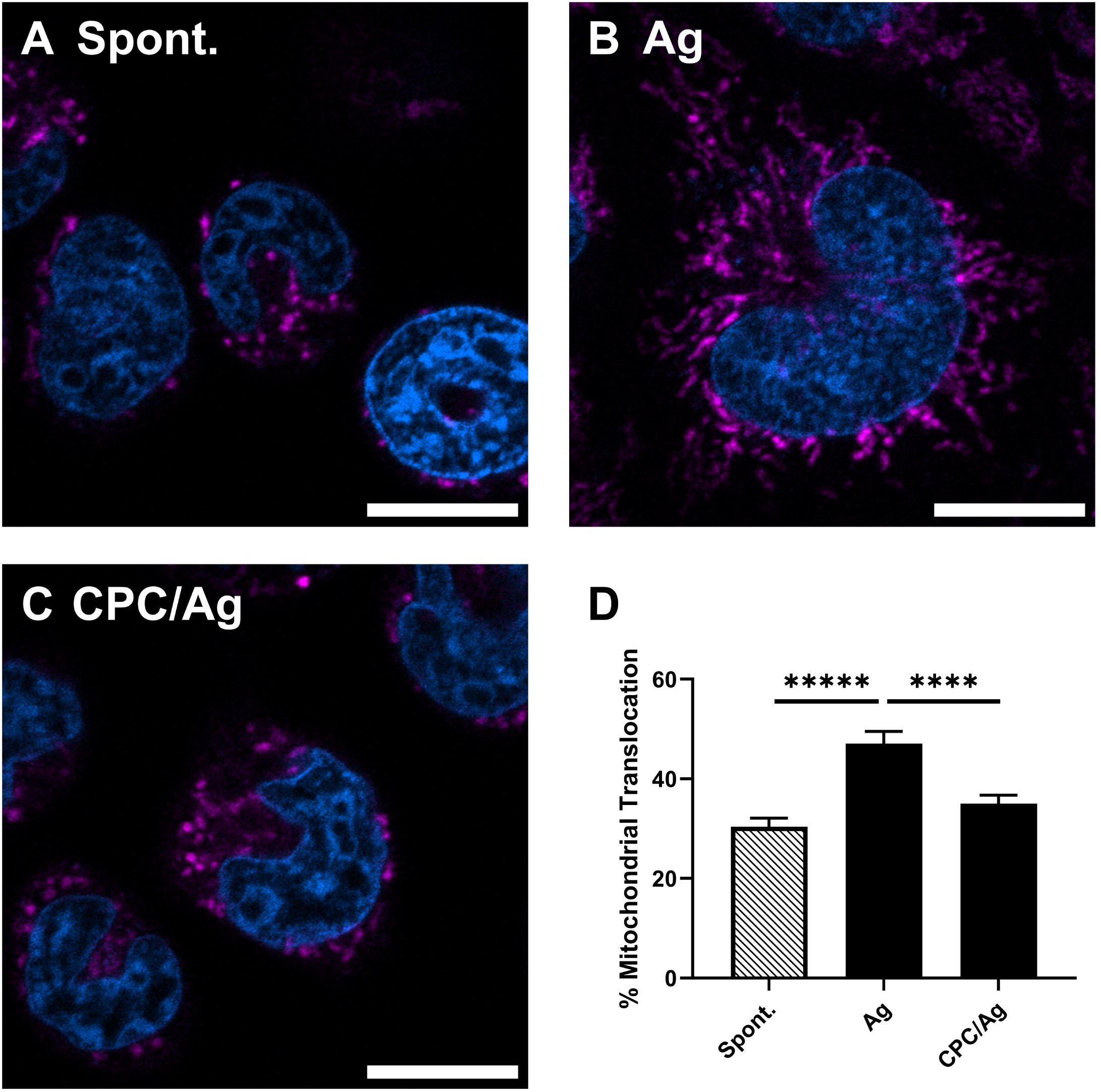
Mitochondrial translocation in antigen-stimulated RBL-2H3 mast cells treated with or without CPC. To assess mitochondrial position, cells were first stained with Hoechst 3342 (nucleus; blue) and Mitotracker FM (mitochondria; magenta) as described in Materials and Methods, then washed and placed into BT with the stated treatments for 45 min, followed by immediate Airyscan imaging of live cells within ∼15 min. Representative images of cells treated with **(A)** only BT (“Spont.”; N = 115 cells), **(B)** 0 µM CPC + antigen (N = 76), or **(C)** 10 µM CPC + antigen (N = 101). Scale bar represents 10 µm. **(D)** Quantification of mitochondrial localization away from the nucleus and toward the plasma membrane. Data are mean ± SEM and represent 3 independent experiments. Statistical significance was determined through one-way ANOVA with Tukey’s post-hoc testing: \*\*\*\**p* < 0.001, \*\*\*\*\**p* < 0.0001.

### CPC causes mitochondrial fission in both Ca^2+^ ionophore- and antigen-stimulated cells

From these mitochondrial translocation images, we also extracted morphological information of the mitochondria in each treatment group. Several morphological features of mitochondria can be used to indicate either mitochondrial fission or fusion. These features include mitochondrial area, perimeter, interconnectivity (area/perimeter), and branching characteristics (Dagda et al. 2009; Wiemerslage and Lee 2016; Chaudhry et al. 2020; Hemel et al. 2021; Chu et al. 2022). Decreased mitochondrial area, perimeter, interconnectivity, and branching characteristics suggest fission, while the inverse would suggest fusion.

We found that Ca^2+^ ionophore alone appears to increase mitochondrial fission (Fig. 4A *versus* 4B), significantly so via the interconnectivity measure (Fig. 6A). Addition of CPC (2.5 µM), in conjunction with Ca^2+^ ionophore stimulation, reduces the mean area, perimeter, interconnectivity, and branch length even further, suggesting that CPC enhances mitochondrial fission (Fig. 6A). These data, taken together with the Drp1 phosphorylation data (Fig. 3), suggest that fission underlies the mechanism for CPC enhancement of mitotranslocation during Ca^2+^-driven mast cell stimulation.

**Figure 6.**
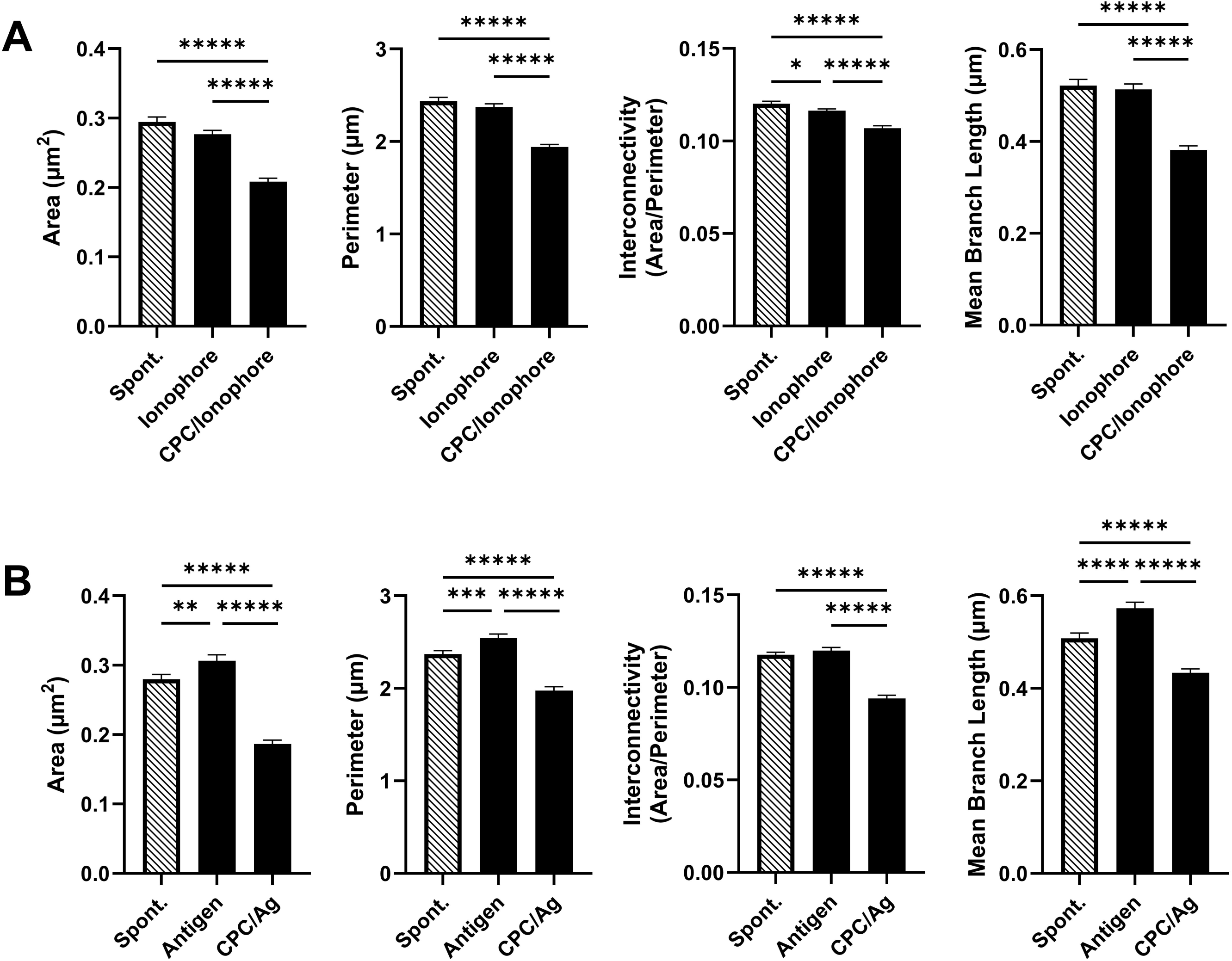
Effects of CPC on mitochondrial structural parameters in RBL-2H3 mast cells, in conjunction with ionophore or antigen stimulation. The calculated area, perimeter, interconnectivity, and mean branch length of mitochondrial networks for each group assessed in the mitochondrial translocation experiments (Figs. 4 & 5). (**A; ionophore stimulation)** Data from cells treated with BT (“Spont.” N = 10,848 mitochondrial elements), 0 µM CPC + 300 nM ionophore (N = 13,692), or 2.5 µM CPC + 300 nM ionophore (N = 15,826). **(B; antigen stimulation)** Data from cells treated with BT (“Spont.” N = 8,097 mitochondrial elements), 0 µM CPC + antigen (N = 14,466), or 2.5 µM CPC + antigen (N = 10,831). Values are mean ± SEM. Statistical significance was determined through one-way ANOVA with Tukey’s post-hoc testing: \**p* < 0.1, \*\**p* < 0.05, \*\*\**p* < 0.01, \*\*\*\**p* < 0.001, \*\*\*\*\**p* < 0.0001. If a comparison is not shown, it was not significant.

In contrast to the ionophore results, antigen stimulation actually increases the mean area, perimeter, and branch length of mitochondria (Fig. 6B and see Fig. 5A *versus* 5B), suggesting antigen-induced fusion of mitochondria. However, concurrent treatment with CPC, along with antigen, decreases the mean area, perimeter, interconnectivity, and branch length significantly below both control and antigen-alone cells (Fig. 6B and Fig. 5B *versus* 5C). In conjunction with their concurrent mast cell stimulants, both 2.5 µM CPC (Fig. 6A) and 10 µM CPC (Fig. 6B) affect these parameters to a similar extent, suggesting that 2.5 µM is sufficient to maximize mitochondrial fission under these conditions. However, these results suggest that CPC-induced mitochondrial fission is not sufficient to overcome CPC-induced inhibition of early events (*e.g*., tyrosine phosphorylation) in the antigen pathway, since CPC-induced fission does not lead to increased mitotranslocation (Fig. 5) or degranulation (Obeng et al. 2023), in the presence of antigen stimulation.

### CPC does not inhibit microtubule polymerization in ionophore-stimulated mast cells

The fact that CPC induces mitochondrial fission in both antigen- and Ca^2+^ ionophore-stimulated cells (Figs. 6A, B), yet only increases mitotranslocation in Ca^2+^-ionophore stimulated cells, raises the question of what factor is contributing to mitochondrial transport to the cellular perimeter in Ca^2+^ ionophore-stimulated cells (Fig. 4) but not in antigen-stimulated cells (Fig. 5). We previously published that microtubule polymerization is inhibited by CPC during antigen stimulation (Obeng et al. 2023), so we aimed to assess CPC effects on microtubules during ionophore stimulation.

To do so, cells were transfected with EGFP-α-tubulin and MitoTracker Red FM to visualize microtubule polymerization simultaneously with mitochondrial movement in RBL-2H3 mast cells (representative cells are shown in Fig. 7). Figure 7A represents a “Spont.” cell with relatively little microtubule polymerization and mitochondria tucked closely beside the nucleus. Figure 7B is a Ca^2+^ ionophore-stimulated mast cell undergoing microtubule polymerization and displaying mitochondrial translocation. Figure 7C displays a representative cell that is both treated with CPC and is Ca^2+^ ionophore-stimulated, showing robust microtubule polymerization and mitochondrial translocation. The majority of ionophore-activated, EGFP-α-tubulin-transfected cells displayed the microtubule polymerization phenotype, whether in the absence or presence of CPC (data not shown).

**Figure 7.**
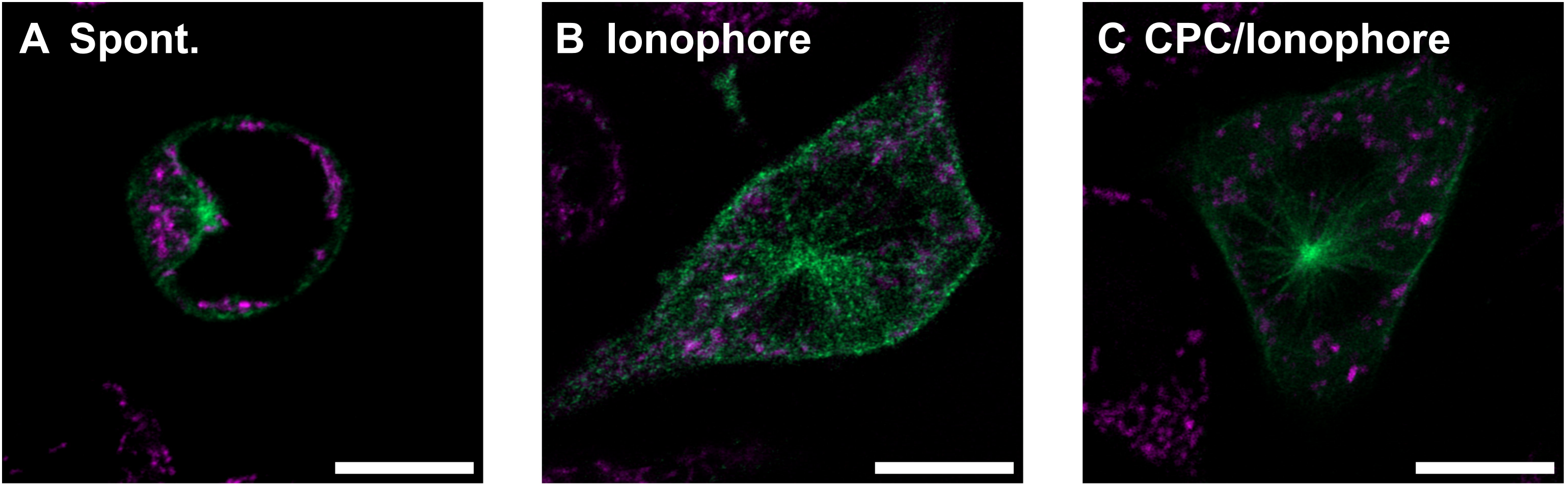
Microtubule polymerization in ionophore-stimulated RBL-2H3 cells with or without CPC treatment. To assess the microtubule polymerization of mast cells, cells were transfected with EGFP--tubulin (microtubules; green). The next day, they were stained with MitoTracker FM (mitochondria; magenta) as described in Materials and Methods, then washed and placed into BT with the stated treatments for 1 hr, followed by immediate Airyscan imaging of live cells within ∼30 min. Representative images of cells treated with **(A)** only BT (“Spont.”; N = 61 cells), **(B)** 300 nM Ca^2+^ ionophore (N = 35) or **(C)** 2.5 µM CPC and 300 nM Ca^2+^ ionophore (N = 52) for 1 hr. Scale bar represents 10 µm.

### CPC differentially impacts mitochondrial membrane potential depending on stimulation type

Mitochondria must maintain sufficient voltage difference across the inner mitochondrial membrane (MMP) to allow for production of ATP (Mitchell 1966). Decrease in MMP due to treatment with a canonical mitochondrial uncoupler attenuates mast cell activation (Mohr and Fewtrell 1987). Thus, we hypothesized that the mitotoxicant CPC may also reduce MMP.

We analyzed the MMP in each treatment group of the mitotranslocation images by measuring the fluorescence intensity of cellular MitoTracker Red FM (Buravkov et al. 2014; Weatherly et al. 2018) in each treatment group. A significant increase in MMP is seen after 1 hr exposure to Ca^2+^ ionophore—surprisingly, further increasing when CPC is added (Fig. 8A). Antigen stimulation also increases MMP, although addition of 10 µM CPC along with the antigen attenuates this response (Fig. 8B). Both stimulation types energize mitochondria, as evidenced by increases in MMP, but CPC produces opposite effects in antigen-vs. ionophore-stimulated cells.

**Figure 8.**
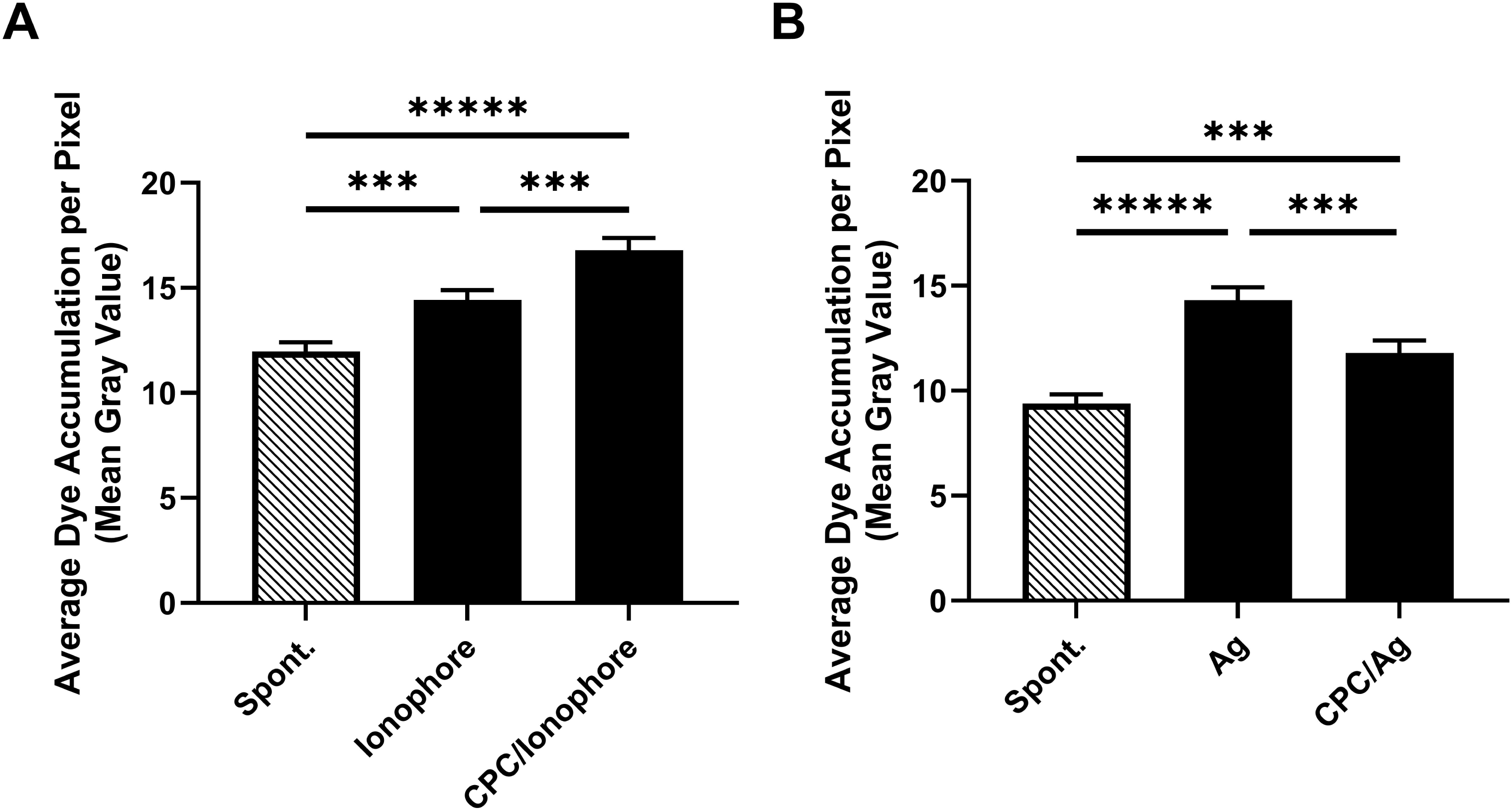
Effects of CPC on mitochondrial membrane potential (MMP) of RBL-2H3 mast cells, in conjunction with Ca^2+^ ionophore or antigen stimulation. The Mitotracker Red dye mean gray value was measured to assess dye accumulation, a proxy for MMP, for each group assessed in the mitochondrial translocation experiments (Figs. 4 & 5). **(A; ionophore stimulation)** Data from cells treated with BT (“Spont.” N = 172 cells), 0 µM CPC + 300 nM ionophore (N = 157), or 2.5 µM CPC + 300 nM ionophore (N = 167). **(B; antigen stimulation)** Data from cells treated with BT (“Spont.” N = 144 cells), 0 µM CPC + antigen (N = 86), or 2.5 µM CPC + antigen (N = 118). Values are mean ± SEM and represent three or four independent experiments. Statistical significance was determined through one-way ANOVA with Tukey’s post-hoc testing: \*\**p* < 0.05, \*\*\**p* < 0.01, \*\*\*\**p* < 0.001, \*\*\*\*\**p* < 0.0001.

## Discussion

In this study, we first set out to fill a gap in toxicological knowledge, regarding the effects of a prominent quaternary ammonium compound on immune cell signaling. We aimed to fully map the biochemical mechanism(s) by which CPC inhibits mast cell signaling and function, to serve as a model for how quaternary ammonium compounds may affect not only mast cells but also related immune cells. Previously, we had determined that CPC targets a few specific proteins’ activation by tyrosine phosphorylation (Obeng et al. 2024) and, separately, the key signaling lipid PIP_2_ (Raut et al. 2022; Aho et al. 2025), but potential CPC effects on immune signaling downstream of Ca^2+^ mobilization remained unknown. *First*, this investigation led to the finding that exposure-relevant doses of CPC do not potently inhibit molecules engaged downstream of Ca^2+^ influx–focusing its mode of action on the aforementioned targets. *Second*, and unexpectedly, CPC causes powerful stimulation of mast cell signaling under inducement of signaling proximal to Ca^2+^ entry, suggesting that immune pathways alternative to the classic IgE-mediated pathway may be activated by this compound. *Third*, we determined the mechanism of this stimulatory effect, which highlights the interrelationship between mitochondrial and immune signaling. *Finally*, these results suggest a novel type of hormetic effect that can flip, depending on cellular energy status.

### CPC does not potently inhibit molecules engaged downstream of Ca^2+^ influx

Employing Ca^2+^ ionophore as an experimental tool allows researchers to determine whether a chemical targets cellular events upstream or downstream of Ca^2+^ influx, or both. Calcium ionophore shuttles Ca^2+^ into the cell, activating enzymes and microtubules critical for degranulation, and bypassing all events before Ca^2+^ influx, including the early tyrosine phosphorylation events and PIP_2_ cleavage (Fig. 1). Because CPC *does not* inhibit mast cell degranulation under Ca^2+^ ionophore stimulation, this result indicates that CPC is not inhibitory toward biochemical targets downstream of the CRAC channel. This finding (Fig. 2) not only reinforces the importance of the tyrosine phosphorylation and PIP_2_ mechanisms of inhibition, but also leads to the question of how CPC actually enhances the process when those are bypassed.

### CPC causes powerful stimulation of Ca^2+^-induced mast cell signaling

Because previous research showed robust CPC inhibition of both human (Obeng et al. 2024) and rat mast cell function, under a wide variety of IgE-mediated FcεRI-activation conditions (Obeng et al. 2023), we were surprised to find that CPC (at non-cytotoxic doses, Fig. S2) actually stimulates Ca^2+^-activated degranulation. For context, arsenite also inhibits mast cells specifically by targeting Syk phosphorylation--but the metalloid displayed no stimulatory (nor inhibitory) effect in conjunction with Ca^2+^ ionophore, showing that its only acute target in the mast cell is upstream (Shim, Kennedy, et al. 2016). This comparison indicates that CPC must initiate an unknown and perhaps unique stimulatory mechanism. Acute exposure with just 1 µM CPC stimulates degranulation by nearly 3-fold, rising to maximal stimulation over control by 2.5 µM CPC–then degranulation falls to baseline levels by 10 µM CPC (Fig. 2).

### Role of cellular energy status suggesting another mode of hormesis

Next, this CPC-induced degranulation stimulation was tested using additional treatment media, first in BT buffer with the glucose swapped for galactose (Fig. S3). Cells were low on electron sources for catabolism due to the fact that galactose is shunted by the cell for other uses such as anabolism (discussed in Weatherly et al. 2018). As expected, the extent of degranulation in the ionophore-only sample (3.5%) was lower than that in the equivalent experiment in the presence of glucose (7%, Fig. 2) – a sign of lower energy status in the Fig. S3 samples. However, the stimulatory effect of CPC on degranulation was still observed, peaking as before at 2.5 µM CPC (Fig. S3).

Next, we assayed the effects of CPC on cells in galactose-DMEM media (the same was used in (Weller et al. 2024)) (Fig. S4), which forces cells to primarily utilize mitochondrial oxidative phosphorylation for energy (described in Weatherly et al. 2016). In this media, CPC causes potent mitochondrial toxicity (ATP, oxygen consumption, and fission (Weller et al. 2024). First, under conditions of very low levels of Ca^2+^ ionophore-stimulated degranulation, CPC exhibited no effect on degranulation (Fig. S4A), in line with its published lack of effect on “spontaneous” degranulation (Obeng et al. 2023). Second, under higher (but still low, 10.8% of granules released) levels of ionophore-stimulated degranulation, CPC enhanced degranulation though the effect did not reach statistical significance (Fig. S4B), potentially because, at the tested doses and in galactose DMEM media, CPC is known to inhibit mitochondrial ATP production (Weller et al. 2024). In contrast, under the stimulatory conditions of Fig. 2, glucose is present: CPC does not drastically inhibit ATP in the presence of glucose (Weller et al. 2024). Thus, while mitochondria undergo fission under these conditions (Weller et al. 2024), ATP is simultaneously inhibited, so cells do not have sufficient energy needed to harness the fissioned mitochondria for degranulation support. There is, still, an echo of the stimulation seen in Fig. 2, likely propelled by residual cellular energy.

As a final energy variation, we analyzed data from ionophore-stimulated degranulation also performed in the galactose-DMEM media but resulting in higher (“moderate,” 20.4%) degranulation in the no-CPC control (Fig. S4C). Here, CPC’s response actually flipped dramatically, causing strong inhibition at CPC doses as low as 0.1 µM. This severe inhibition may be explained by the known, strong inhibition of mitochondrial electron flow (Ledue et al. 2026) and ATP production due to CPC when in galactose DMEM (Weller et al. 2024). Therefore, in the galactose-DMEM, in which the only pathway cells have to create energy is potently inhibited by CPC, mast cells are unable to meet the energy demands needed for degranulation when higher (*e.g.*, Fig. S4C) levels of ionophore-induced cell stimulation happen. This potent mitochondrial ATP inhibition then outweighs the stimulatory effect of CPC-induced fission-supported mitotranslocation. The inhibition in Fig. S4C may also be due in part to the PIP_2_ inhibition as well because PIP_2_ is also involved in the SNARE exocytotic machinery (Dai et al. 2007).

Both the initial U-shaped stimulation curve (Fig. 2, Fig. S3) and also the flipping of CPC’s effect on immune signaling due to different modes of stimulation and when in different metabolic states (Fig. S4) reminds the authors of hormesis, involving stimulation of a physiological process at lower doses while inhibition occurs at higher doses (Mattson 2008). Further, we believe the energy-dependent “flipping” from stimulation back to potent inhibition may be a novel dimension of hormesis, with some previous literature evidence (Zhang et al. 2008). In real physiological conditions, tissues may vacillate, flipping depending on energy demands and glucose availability. Also, we previously showed that IgE-mediated degranulation is inhibited (and not, under any condition, stimulated) by CPC to the same degree, whether the cells are in galactose or in glucose (Weller et al. 2024). Whether CPC effects on degranulation are stimulatory/inhibitory is a function of stimulation method.

### Alternative, non-IgE-mediated, modes of mast cell stimulation

Besides IgE receptor-mediated signaling, there exist in nature multiple alternative pathways for stimulation of mast cell degranulation which bypass (CPC-inhibited) early phosphorylation events and thus may be prone to stimulation by CPC. Estradiol and some environmental estrogens (Narita et al. 2007; Zaitsu et al. 2007; Kennedy et al. 2012) initiate mast cell degranulation through estrogen receptor alpha in a manner involving Ca^2+^ influx from the external medium (Zaitsu et al. 2007) which may involve activation of alternative Ca^2+^ channels (Gui et al. 2025). Separately, a wide variety of ligands (e.g., endothelin-1, neurotensin, Substance P, various basic secretagogues) have been shown to activate various G protein-coupled receptors on the surface of mast cells in a manner that induces degranulation via a mechanism proximal to SOCE (Redegeld et al. 2018). For example, Mas-related G protein-coupled receptors like MRGPRX2 bind ligands, transmitting their signal to G proteins which undergo nucleotide exchange, thereby stimulating phospholipase Cβ (Ferry et al. 2001), which cleaves PIP_2_, swiftly initiating SOCE and, thus, degranulation–without the involvement of early tyrosine phosphorylation events (Tatemoto et al. 2006; Subramanian et al. 2016). Since MRGPRX2 signaling does not rely on Syk/LAT signaling (inhibited targets of CPC), CPC may stimulate degranulation in this system as with Ca^2+^ ionophore (if CPC stimulation of mitotranslocation outweighs its inhibition of PIP_2_). (Note: evidence of a Mas-related G protein-coupled receptor Mrgprb3 is found in RBL-2H3 cells but is non-responsive (Sahid et al. 2020); also there is no direct CPC stimulation of it, evidenced by the lack of CPC-alone stimulation of RBL-2H3 degranulation (Obeng et al. 2023). Another potential stimulation mechanism that skirts both early phosphorylation and PIP_2_ is SOCE induction via inhibition of the sacro/endoplasmic reticulum Ca^2+^ ATPase pump on the endoplasmic reticulum by the botanical medicine thapsigargin (Lytton et al. 1991; Andersen et al. 2015; Shim, Kennedy, et al. 2016). Additionally, the Toll-like receptor 2 can be activated by the LL37 peptide in rosacea (Fan et al. 2025), using JAK/STAT activation to induce degranulation. Overall, the possibilities for mast cell activation via pathways that may be enhanced by CPC are many.

### Mechanism underlying CPC’s stimulatory effect: role of mitochondrial fission

Previous researchers discovered that Substance P-activated mast cell degranulation is supported by Drp1-driven mitotranslocation (Zhang et al. 2011; Zhang et al. 2012), and inhibition of Drp1 inhibits degranulation (Wang et al. 2022). Because CPC induces mitochondrial fission (Weller et al. 2024), we hypothesized that this fission underlies CPC’s enhancement of ionophore-driven degranulation. Because Drp1 is a driver of mitochondrial fission (Zhang et al. 2011; Simula et al. 2018; Kamerkar et al. 2025) and it is activated by phosphorylation at its residue Ser616 (Kamerkar et al. 2025), we assayed CPC effects on Drp1 Ser616 and found its phosphorylation is increased in RBL-2H3 mast cells (Fig. 3). (It was previously shown that Drp1 Ser616 phosphorylation does not change in response to IgE-mediated mast cell activation (Pavlyuchenkova et al. 2022). While authors (Wang et al. 2022) did not explicitly connect Drp1-driven mitochondrial fission to mitotranslocation and thus to Ca^2+^-proximal activation of mast cells, theirs and our results align with this hypothesis.

With the Drp1 result as a hypothesis driver, we next directly assessed the level of mitochondrial fusion/networking versus fission. We followed mitochondria, using Mitotracker Red FM (mitochondrial stain) and Hoechst 33342 (nuclear stain) and Airyscan microscopy. The superior resolution of Airyscan (∼1.7-fold improvement over the ∼215 nm calculated for Mitotracker Red FM and the microscope setup), compared to conventional confocal microscopy, enables detailed analysis of mitochondrial morphology, including area, perimeter, interconnectivity, and branching of mitochondria. For example, some highly significant effects displayed in Fig. 6 involve ∼100 nm changes, which would be blurred in traditional confocal. Ca^2+^ ionophore alone modestly increases mitochondrial fission (Fig. 4A *versus* 4B; Fig. 6A), and addition of CPC further increases measures of fission (Fig. 6A). These experiments were conducted in glucose, which protects cells from losing ATP production (Weller et al. 2024), but CPC-induced fission still occurs whether in glucose (Fig. 6) or galactose (Weller et al. 2024) – an effect also described with proton ionophore mitotoxicant triclosan (Weatherly et al. 2018). These data, taken together with Drp1 phosphorylation (Fig. 3), suggest that mitochondrial fission underlies the mechanism for CPC enhancement of mitotranslocation and thus degranulation during Ca^2+^-driven mast cell stimulation.

In contrast to the ionophore results, IgE-mediated stimulation actually increases multiple parameters related to mitochondrial fusion (Fig. 6B and see Fig. 5A *versus* 5B)--but addition of CPC causes mitochondrial fission even upon this IgE-mediated background (Fig. 6B and Fig. 5B *versus* 5C). Overall though, the IgE-mediated results indicate that CPC-induced mitochondrial fission is insufficient for overcoming CPC-induced inhibition of early signaling events (Fig. 5) (Obeng et al. 2023).

### Mechanism underlying CPC’s stimulatory effect: role of mitochondrial translocation and microtubules

With this confirmation that CPC indeed induces mitochondrial fission in the ionophore-treated cells, we investigated whether CPC also affects mitotranslocation. For these experiments, we utilized a 2.5 µM CPC dose since it produced the maximal amount of stimulation in conjunction with Ca^2+^ ionophore (Fig. 1). Ca^2+^ ionophore increases mitochondrial translocation, an effect that is enhanced upon concurrent addition of CPC (Fig. 4). These data suggest that increased mitochondrial fission and, thus, translocation, possibly to provide localized mitochondrial ATP generation for granule exocytosis (Zhang et al. 2011), are a likely mechanism for the increase in ionophore-initiated degranulation caused by CPC. There exists little information in the literature regarding chemical stimulation of mitochondrial translocation, but cancer tissues can enhance the process (Agarwal et al. 2019).

In contrast, while antigen treatment significantly increased mitochondrial translocation, concurrent CPC exposure decreased translocation, back down to control (Fig. 5). CPC causes approximately the same amount of mitochondrial fission in both antigen- and Ca^2+^ ionophore-stimulated groups (Fig. 6), yet increases mitotranslocation only in the Ca^2+^ ionophore-stimulated cell.

This difference between CPC responses upon IgE-mediated- and Ca^2+^ ionophore-stimulation (Fig. 4D vs. 5D) raises the question of how mitochondria in ionophore-stimulated cells are transported to the cellular periphery. We previously showed that CPC inhibits IgE-mediated microtubule polymerization (Obeng et al. 2023), so we hypothesized that the opposite occurs in the presence of CPC & ionophore. Our results demonstrate that microtubules (green) do still polymerize in Ca^2+^ ionophore-stimulated mast cells treated with CPC (Fig. 7), allowing mitochondria (magenta) to be transported to the cellular periphery. Thus, microtubule polymerization is crucial to the mechanism of mitotranslocation-enhanced degranulation.

### Mechanism underlying CPC’s stimulatory effect: role of mitochondrial membrane potential and electron flow

To further understand the mitochondrial mechanism underlying CPC stimulation of Ca^2+^ ionophore-induced degranulation, we analyzed MMP. Generated by the mitochondrial electrochemical gradient, MMP is crucial to enable ATP synthesis (Zorova et al. 2018; Piotin et al. 2024). Both stimulation methods tested were associated with enhancement of MMP: mitochondria become energized in mast cell stimulation.

Interestingly, CPC further enhanced Ca^2+^ ionophore-boosted MMP (Fig. 8A). In contrast, CPC decreased antigen-stimulated MMP, back to control levels (Fig. 8B). Because the 10 µM CPC in Fig. 8B antigen-stimulated cells reduces MMP, this result may represent a tipping point at which higher [CPC] reduce mitochondrial energy in the mast cell signaling context. Overall, the MMP increase due to CPC/Ca^2+^ ionophore suggests energized mitochondria under the conditions showing stimulated degranulation.

In order to understand these MMP effects, we next assayed electron flow through Complex I of the electron transport chain (Fig. S9). Electron flow through the mitochondrial respiratory chain powers proton pumping, maintaining MMP. As we previously published (Weller et al. 2024; Ledue et al. 2026), CPC inhibits electron flow, in both quiescent and Ca^2+^ ionophore-stimulated mast cells (Fig. S9A). Separately, ionophore stimulation of the mast cells increased electron flow, whether ± CPC– resulting in the same stimulated degranulation as in Fig. 2 (Fig. S9B). This increased electron flow may partially explain the increased MMP (though cannot explain the Fig. 8A results fully due to CPC inhibition of NADH oxidation). Ionophore increases electron flow (Fig. S9) and, thus, MMP (Fig. 8A). Ionophore-induced increase in cytosolic [Ca^2+^] may activate the mitochondrial Ca^2+^ uniporter, thereby increasing [Ca^2+^] in the matrix, thus activating catabolic enzymes (Denton et al. 1978; Lawlis and Roche 1981). While CPC causes Ca^2+^ efflux from the mitochondria in galactose media-bathed cells undergoing mitochondrial ATP inhibition (Weller et al. 2024), there is no such CPC-induced Ca^2+^ efflux to counter the ionophore-induced Ca^2+^ influx under glucose conditions (Weller et al. 2024), as in the current study.

### Relevance to other immune cell types and to vaccine adjuvant function

The CPC mechanisms described in this manuscript may also extend to other immune cells, such as T cells, which share common signaling elements with mast cells. For example, mitochondria also move to the polarized leading edge of migrating T cells (Campello et al. 2006) and also translocate, in a manner crucial for activation, toward the immunological synapse formed between T helper and antigen presenting cells (Quintana et al. 2007).

Interestingly, vaccine developers have recently begun using CPC as an immunological adjuvant in both mice (Larsen et al. 2004) and human clinical trials (Deming et al. 2025) for intranasal vaccines. One postulated mechanism is that strong CPC interaction with mucin significantly increases nasal retention of the associated antigenic payload, enhancing immunogenicity (Wong et al. 2014). However, another mechanism of CPC efficacy as an adjuvant could potentially be CPC induction of mitochondrial fission and thus translocation, activating immune cells.

### Limitations, future work, and conclusions

A limitation of this study is that CPC induction of mitochondrial fission and translocation may not be the sole mechanism accounting for the entirety of CPC’s stimulatory effect on mast cell degranulation. The degree to which mitochondrial translocation modulates degranulation has not yet been quantified, to our knowledge, an area ripe for investigation. Other avenues of future research include quaternary ammonium compound effects on other modes of mast cell activation and immune cell types. The importance of mast cell function in the brain (Silver and Curley 2013), the ability of environmentally-relevant CPC exposures to accumulate in the brain (Pottel et al. 2020; Cohn et al. 2024), and the critical nature of mitochondria in energy-intensive brain function point to future investigations. With a growing body of evidence on the mechanistic effects of CPC, additional targeted pharmacokinetic and epidemiological studies are warranted.

In this study, we have pinpointed the inhibitory mechanism of CPC to the upstream portion of IgE-mediated mast cell signaling and elucidated a stimulatory mechanism of CPC on Ca^2+^ ionophore-mediated mast cell degranulation (Fig. 9). This finding solidifies the conclusion that CPC inhibition of mast cell signaling is targeted to events (*e.g.* Syk tyrosine phosphorylation, PIP_2_) upstream of Ca^2+^ influx. This stimulatory mechanism is characterized by CPC-induced mitochondrial fission, energization (increased MMP), and translocation mediated by microtubules–and may be applicable to other Ca^2+^-proximal mast cell signaling pathways and immune cell types. However, CPC’s inhibition of MMP and mitotranslocation in the IgE-mediated background demonstrates that induction of fission alone is not sufficient to boost degranulation. We have also demonstrated that stimulatory effects of CPC may depend on cellular energy status, flipping CPC stimulation to inhibition. This research not only connects mitochondrial and immune cell signaling to inform basic cell biology, but also further reveals mechanisms of CPC immune modulation. These findings may also inform our understanding of the impacts of other quaternary ammonium compounds on immune cells, ultimately helping consumers and regulators make informed decisions about these drugs.

**Figure 9.**
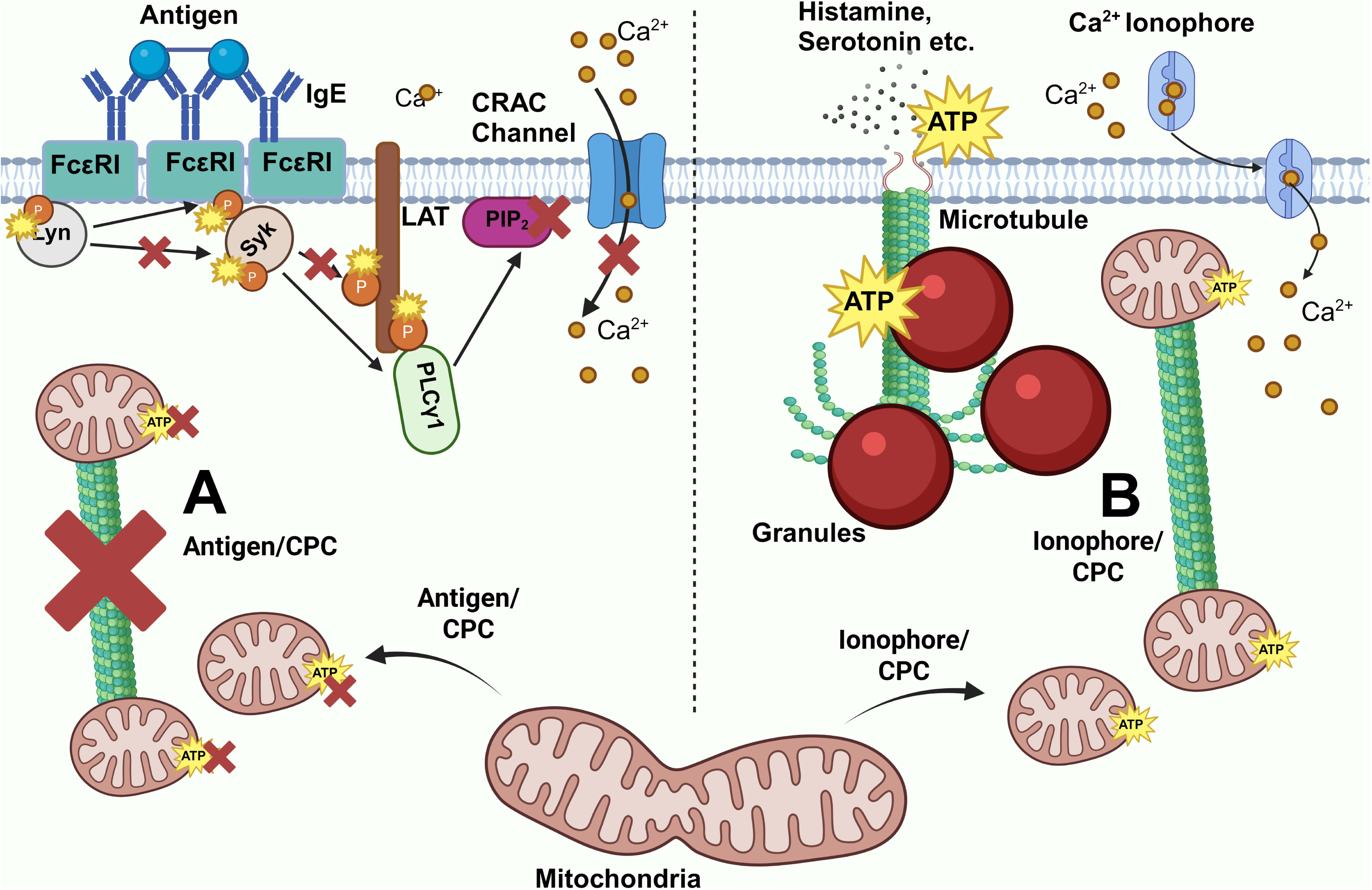
A graphical summary of CPC effects on mitotranslocation under either antigen and Ca^2+^ ionophore stimulation of mast cells. **(A)** Under antigen stimulation, CPC decreases mitotranslocation (red X indicates inhibition) despite increasing mitochondrial fission (arrow pointing away indicates fission), indicating that CPC-induced mitochondrial fission may be necessary but not sufficient for mitotranslocation and degranulation. In antigen-stimulated mast cells, this phenomenon is likely caused by CPC inhibition of microtubule polymerization due to inhibition of tyrosine phosphorylation and thus SOCE. **(B)** Under Ca^2+^ ionophore stimulation, CPC increases mitotranslocation via increased mitochondrial fission along with robust microtubule polymerization, leading to enhanced degranulation – indicating that CPC-induced mitochondrial fission may be sufficient to cause enhanced degranulation in certain immune signaling pathways. This finding solidifies the conclusion that CPC inhibition of mast cell signaling is targeted to events upstream of Ca^2+^ influx.

## Supplementary material

Supplementary material is available at *Toxicological Sciences* online.

## Acknowledgments

We thank Marisa Johnson, Isaac Dostie, Noah Adelman, Ellis Klick, Quinn D’Alessio, and Miriam Klee for lab support and discussions. We thank Dr. Melissa Maginnis for allowing us to use the NanoDrop machine and lab equipment obtained through the Maine IDeA Network of Biomedical Research Excellence program under NIH-National Institute of General Medical Sciences grant number P20GM0103423. The authors would like to acknowledge the Microscopy and Image Analysis Core facility (RRID:SCR_025784), supported by NIH award number 1P20GM144265, for use of Zeiss LSM980 Confocal Microscope/Imaris software on a fee for service basis and Dr. Zhengxin Ma for training, discussion, and help collecting the data shown in Figures 4, 5, and 7.

## References

Abezgauz L et al. 2010. Effect of Hofmeister anions on micellization and micellar growth of the surfactant cetylpyridinium chloride. J Colloid Interface Sci. 342(1):83–92. 10.1016/j.jcis.2009.08.045

Abramson J, Pecht I. 2007. Regulation of the mast cell response to the type 1 Fcε receptor. Immunological Reviews. 217(1):231–254. 10.1111/j.1600-065X.2007.00518.x

Agarwal E et al. 2019. Myc Regulation of a Mitochondrial Trafficking Network Mediates Tumor Cell Invasion and Metastasis. Molecular and Cellular Biology. 39(14):e00109–19. 10.1128/MCB.00109-19

Aho BM, Wagner DJ, Gosse JA, Hess ST. 2025. Nanoscale clustering and dynamics of phosphatidylinositol 4,5-bisphosphate in an immune cell model. Biophysical Journal. 0(0). https://www.cell.com/biophysj/abstract/S0006-3495(25)00423-0. 10.1016/j.bpj.2025.07.004

Alsaleh NB, Persaud I, Brown JM. 2016. Silver Nanoparticle-Directed Mast Cell Degranulation Is Mediated through Calcium and PI3K Signaling Independent of the High Affinity IgE Receptor. PLoS One. 11(12):e0167366. 10.1371/journal.pone.0167366

Andersen TB et al. 2015. Thapsigargin—From Thapsia L. to Mipsagargin. Molecules. 20(4):6113–6127. 10.3390/molecules20046113

Arnold WA et al. 2023. Quaternary Ammonium Compounds: A Chemical Class of Emerging Concern. Environ Sci Technol. 57(20):7645–7665. 10.1021/acs.est.2c08244

Bernauer U et al. 2015. Opinion on Cetylpyridinium chloride -Submission II: COLIPA n° P97. https://data.europa.eu/doi/10.2875/962022

Berridge MJ. 1993. Inositol trisphosphate and calcium signalling. Nature. 361(6410):315–325. 10.1038/361315a0

Biomonitoring California. 2026. Biomonitoring California Priority Chemicals. California Government. https://biomonitoring.ca.gov/chemicals/priority-chemicals

Bonesvoll P, Gjermo P. 1978. A comparision between chlorhexidine and some quaternary ammonium compounds with regard to retention, salivary concentration and plaque-inhibiting effect in the human mouth after mouth rinses. Arch Oral Biol. 23(4):289–294. 10.1016/0003-9969(78)90021-3

Buravkov SV, Pogodina MV, Buravkova LB. 2014. Comparison of mitochondrial fluorescent dyes in stromal cells. Bull Exp Biol Med. 157(5):654–658. 10.1007/s10517-014-2637-3

Campello S et al. 2006. Orchestration of lymphocyte chemotaxis by mitochondrial dynamics. J Exp Med. 203(13):2879–2886. 10.1084/jem.20061877

Chaudhry A, Shi R, Luciani DS. 2020. A pipeline for multidimensional confocal analysis of mitochondrial morphology, function, and dynamics in pancreatic β-cells. American Journal of Physiology-Endocrinology and Metabolism. 318(2):E87–E101. 10.1152/ajpendo.00457.2019

Chávez E, Bravo C. 1982. Anisotropic action of cetyl pyridinium chloride on rat heart mitochondria. Archives of Biochemistry and Biophysics. 213(1):81–86. 10.1016/0003-9861(82)90442-8

Chu C-H, Tseng W-W, Hsu C-M, Wei A-C. 2022. Image Analysis of the Mitochondrial Network Morphology With Applications in Cancer Research. Front Phys. 10. https://www.frontiersin.org/journals/physics/articles/10.3389/fphy.2022.855775/full. 10.3389/fphy.2022.855775

Cohn EF et al. 2024. Pervasive environmental chemicals impair oligodendrocyte development. Nat Neurosci. 1–10. 10.1038/s41593-024-01599-2

Dagda RK et al. 2009. Loss of PINK1 function promotes mitophagy through effects on oxidative stress and mitochondrial fission. J Biol Chem. 284(20):13843–13855. 10.1074/jbc.M808515200

Dai H, Shen N, Araç D, Rizo J. 2007. A Quaternary SNARE-Synaptotagmin-Ca2+-Phospholipid Complex in Neurotransmitter Release. J Mol Biol. 367(3):848–863. 10.1016/j.jmb.2007.01.040

Datta S et al. 2017. In Vitro Evaluation of Mitochondrial Function and Estrogen Signaling in Cell Lines Exposed to the Antiseptic Cetylpyridinium Chloride. Environ Health Perspect. 125(8):087015. 10.1289/EHP1404

Deming ME et al. 2025. An intranasal adjuvanted, recombinant influenza A/H5 vaccine primes against diverse H5N1 clades: a phase I trial. Nat Commun. 16(1):9321. 10.1038/s41467-025-64686-3

Denton RM, Richards DA, Chin JG. 1978. Calcium ions and the regulation of NAD+-linked isocitrate dehydrogenase from the mitochondria of rat heart and other tissues. Biochem J. 176(3):899–906. 10.1042/bj1760899

Environmental Working Group. EWG/Commonweal Study #1, industrial chemicals and pesticides in adults. https://www.ewg.org/sites/humantoxome/participants/participant-group.php?group=bb1

Fan H et al. 2025. LL37-driven mast cell degranulation and inflammation in rosacea via TLR2/JAK2/STAT3 axis. Front Immunol. 16: 1672021. https://www.frontiersin.org/journals/immunology/articles/10.3389/fimmu.2025.1672021/full. 10.3389/fimmu.2025.1672021

Ferry X, Eichwald V, Daeffler L, Landry Y. 2001. Activation of βγ Subunits of Gi2 and Gi3 Proteins by Basic Secretagogues Induces Exocytosis Through Phospholipase Cβ and Arachidonate Release Through Phospholipase Cγ in Mast Cells. J Immunol. 167(9):4805–4813. 10.4049/jimmunol.167.9.4805

Foreman JC, Mongar JL, Gomperts BD. 1973. Calcium Ionophores and Movement of Calcium Ions following the Physiological Stimulus to a Secretory Process. Nature. 245(5423):249–251. 10.1038/245249a0

Frantz AL. 2023. Chronic quaternary ammonium compound exposure during the COVID-19 pandemic and the impact on human health. Toxicol Environ Health Sci. 15(3):199–206. 10.1007/s13530-023-00173-w

Galli SJ et al. 2005. Mast cells as “tunable” effector and immunoregulatory cells: recent advances. Annu Rev Immunol. 23:749–786. 10.1146/annurev.immunol.21.120601.141025

Gauld SB, Dal Porto JM, Cambier JC. 2002. B Cell Antigen Receptor Signaling: Roles in Cell Development and Disease. Science. 296(5573):1641–1642. 10.1126/science.1071546

Gottlieb S. 2019. Safety and Effectiveness of Consumer Antiseptic Rubs; Topical Antimicrobial Drug Products for Over-the-Counter Human Use. https://www.federalregister.gov/d/2019-06791

Gui Z et al. 2025. The role of estrogen receptors in intracellular estrogen signaling pathways, an overview. The Journal of Steroid Biochemistry and Molecular Biology. 245:106632. 10.1016/j.jsbmb.2024.106632

Hemel IMGM, Engelen BPH, Luber N, Gerards M. 2021. A hitchhiker’s guide to mitochondrial quantification. Mitochondrion. 59:216–224. 10.1016/j.mito.2021.06.005

Hrubec TC et al. 2021. Altered toxicological endpoints in humans from common quaternary ammonium compound disinfectant exposure. Toxicol Rep. 8:646–656. 10.1016/j.toxrep.2021.03.006

Hutchinson LM et al. 2011. Inorganic arsenite inhibits IgE receptor-mediated degranulation of mast cells. J Appl Toxicol. 31(3):231–241. 10.1002/jat.1585

Kagayama M, Douglas WW. 1974. ELECTRON MICROSCOPE EVIDENCE OF CALCIUM-INDUCED EXOCYTOSIS IN MAST CELLS TREATED WITH 48/80 OR THE IONOPHORES A-23187 AND X-537A. J Cell Biol. 62(2):519–526. 10.1083/jcb.62.2.519

Kamerkar SC, Liu A, Higgs HN. 2025. Mitochondrial fission – changing perspectives for future progress. J Cell Sci. 138(9):jcs263640. 10.1242/jcs.263640

Kennedy RH et al. 2012. Estrogen Mimetic 4-tert-Octylphenol Enhances Ige-Mediated Degranulation of Rbl-2H3 Mast Cells. Journal of Toxicology and Environmental Health, Part A. 75(24):1451–1455. 10.1080/15287394.2012.722184

Kinet JP. 1999. The high-affinity IgE receptor (Fc epsilon RI): from physiology to pathology. Annu Rev Immunol. 17:931–972. 10.1146/annurev.immunol.17.1.931

Klaassen, C.D.E., Casarett, L.J.T., 2019. Casarett and Doull’s Toxicology.

Kuby, J. 1997. Immunology. Sixth. W.H. Freeman, New York.

Kux L. 2016. Safety and Effectiveness of Consumer Antiseptics; Topical Antimicrobial Drug Products for Over-the-Counter Human Use; Proposed Amendment of the Tentative Final Monograph; Reopening of Administrative Record. https://www.federalregister.gov/d/2016-15410

Larsen ST et al. 2004. Adjuvant effect of quaternary ammonium compounds in a murine model. Toxicology Letters. 151(2):389–398. 10.1016/j.toxlet.2004.03.008

Lawlis VB, Roche TE. 1981. Regulation of bovine kidney alpha-ketoglutarate dehydrogenase complex by calcium ion and adenine nucleotides. Effects on S0.5 for alpha-ketoglutarate. Biochemistry. 20(9):2512–2518. 10.1021/bi00512a023

Ledue EL et al. 2026. Antimicrobial cetylpyridinium chloride disrupts the mitochondrial electron transport chain with potency comparable to cyanide, via cardiolipin interference at cytochrome c. Toxicol Sci. kfag155. 10.1093/toxsci/kfag155

Liu R et al. 2023. Maternal exposure to cetylpyridinium chloride impairs oogenesis by causing mitochondria disorder in neonates. Environ Toxicol Pharmacol. 102:104239. 10.1016/j.etap.2023.104239

Lytton J, Westlin M, Hanley MR. 1991. Thapsigargin inhibits the sarcoplasmic or endoplasmic reticulum Ca-ATPase family of calcium pumps. J Biol Chem. 266(26):17067–17071

Mandal AB, Nair BU. 1991. Cyclic voltammetric technique for the determination of the critical micelle concentration of surfactants, self-diffusion coefficient of micelles, and partition coefficient of an electrochemical probe. J Phys Chem. 95(22):9008–9013. 10.1021/j100175a106

Marano N et al. 1993. Fc RI and the T cell receptor for antigen activate similar signalling pathways in T cell-RBL cell hybrids. Cellular Signalling. 5(2):155–167. 10.1016/0898-6568(93)90067-V

Mattson MP. 2008. Hormesis Defined. Ageing Res Rev. 7(1):1–7. 10.1016/j.arr.2007.08.007

Metcalfe DD, Baram D, Mekori YA. 1997. Mast cells. Physiol Rev. 77(4):1033–1079. 10.1152/physrev.1997.77.4.1033

Metzger H et al. 1982. Structure of the high-affinity mast cell receptor for IgE. Fed Proc. 41(1):8–11

Mitchell P. 1966. Chemiosmotic Coupling in Oxidative and Photosynthetic Phosphorylation. Biological Reviews. 41(3):445–501. 10.1111/j.1469-185X.1966.tb01501.x

Mohapatra S et al. 2023. Quaternary ammonium compounds of emerging concern: Classification, occurrence, fate, toxicity and antimicrobial resistance. Journal of Hazardous Materials. 445:130393. 10.1016/j.jhazmat.2022.130393

Mohr FC, Fewtrell C. 1987. The relative contributions of extracellular and intracellular calcium to secretion from tumor mast cells. Multiple effects of the proton ionophore carbonyl cyanide m-chlorophenylhydrazone. Journal of Biological Chemistry. 262(22):10638–10643. 10.1016/S0021-9258(18)61011-7

Mukerjee P, Mysels K. Critical micelle concentrations of aqueous surfactant systems. National Institute of Standards and Technology. 10.6028/NBS.NSRDS.36

Narita S et al. 2007. Environmental Estrogens Induce Mast Cell Degranulation and Enhance IgE-Mediated Release of Allergic Mediators. Environ Health Perspect. 115(1):48–52. 10.1289/ehp.9378

Nishida K et al. 2005. FcεRI-mediated mast cell degranulation requires calcium-independent microtubule-dependent translocation of granules to the plasma membrane. J Cell Biol. 170(1):115–126. 10.1083/jcb.200501111

Obeng B et al. 2023. Pharmaceutical agent cetylpyridinium chloride inhibits immune mast cell function by interfering with calcium mobilization. Food and Chemical Toxicology. 179:113980. 10.1016/j.fct.2023.113980

Obeng B et al. 2024. Anti-microbial cetylpyridinium chloride suppresses mast cell function by targeting tyrosine phosphorylation of Syk kinase. J Immunotoxicol. 21(1):2443397. 10.1080/1547691X.2024.2443397

Pavlyuchenkova AN et al. 2022. Mitochondria-targeted triphenylphosphonium-based compounds inhibit FcεRI-dependent degranulation of mast cells by preventing mitochondrial dysfunction through Erk1/2. Life Sciences. 288:120174. 10.1016/j.lfs.2021.120174

Piotin A et al. 2024. Oxidative Stress and Mitochondria Are Involved in Anaphylaxis and Mast Cell Degranulation: A Systematic Review. Antioxidants. 13(8):920. 10.3390/antiox13080920

Pottel J et al. 2020. The activities of drug inactive ingredients on biological targets. Science. 369(6502):403–413. 10.1126/science.aaz9906

Putney JW. 1986. A model for receptor-regulated calcium entry. Cell Calcium. 7(1):1–12. 10.1016/0143-4160(86)90026-6

Quintana A et al. 2007. T cell activation requires mitochondrial translocation to the immunological synapse. Proc Natl Acad Sci U S A. 104(36):14418–14423. 10.1073/pnas.0703126104

Raps S et al. 2024. Triclosan and its alternatives, especially chlorhexidine, modulate macrophage immune response with distinct modes of action. Science of The Total Environment. 914:169650. 10.1016/j.scitotenv.2023.169650

Raut P et al. 2022. Cetylpyridinium chloride (CPC) reduces zebrafish mortality from influenza infection: Super-resolution microscopy reveals CPC interference with multiple protein interactions with phosphatidylinositol 4,5-bisphosphate in immune function. Toxicol Appl Pharmacol. 440:115913. 10.1016/j.taap.2022.115913

Redegeld FA et al. 2018. Non-IgE mediated mast cell activation. Immunological Reviews. 282(1):87–113. 10.1111/imr.12629

Rusan NM et al. 2001. Cell Cycle-Dependent Changes in Microtubule Dynamics in Living Cells Expressing Green Fluorescent Protein-α Tubulin182. Molecular Biology of the Cell. 12(4):971–980. 10.1091/mbc.12.4.971

Sahid MNA, Liu S, Mogi M, Maeyama K. 2020. Tachykinin-1 receptor antagonism suppresses substance-P- and compound 48/80-induced mast cell activation from rat mast cells expressing functional mas-related GPCR B3. Inflamm Res. 69(3):289–298. 10.1007/s00011-020-01319-z

Saladino A, Hawkins H, Trump B. 1971. Ion movements in cell injury. Effects of the cationic detergent cetyl pyridinium chloride on the ultrastructure and function of the toad bladder. The American journal of pathology. 64(2): 271–294. https://pubmed.ncbi.nlm.nih.gov/4946878/

Seldin DC et al. 1985. Homology of the rat basophilic leukemia cell and the rat mucosal mast cell. Proc Natl Acad Sci USA. 82(11):3871–3875

Shi Y, Luo HQ, Li NB. 2011. Determination of the critical premicelle concentration, first critical micelle concentration and second critical micelle concentration of surfactants by resonance Rayleigh scattering method without any probe. Spectrochimica Acta Part A: Molecular and Biomolecular Spectroscopy. 78(5):1403–1407. 10.1016/j.saa.2011.01.018

Shim J, Kennedy RH, et al. 2016. Arsenic inhibits mast cell degranulation via suppression of early tyrosine phosphorylation events. Journal of Applied Toxicology. 36(11):1446–1459. 10.1002/jat.3300

Shim J, Weatherly LM, et al. 2016. Triclosan is a Mitochondrial Uncoupler in Live Zebrafish. J Appl Toxicol. 36(12):1662–1667. 10.1002/jat.3311

Silver R, Curley JP. 2013. Mast cells on the mind: new insights and opportunities. Trends Neurosci. 36(9):513–521. 10.1016/j.tins.2013.06.001

Simula L et al. 2018. Drp1 Controls Effective T Cell Immune-Surveillance by Regulating T Cell Migration, Proliferation, and cMyc-Dependent Metabolic Reprogramming. Cell Rep. 25(11):3059–3073.e10. 10.1016/j.celrep.2018.11.018

Smith-Garvin JE, Koretzky GA, Jordan MS. 2009. T Cell Activation. Annu Rev Immunol. 27:591–619. 10.1146/annurev.immunol.021908.132706

Subramanian H, Gupta K, Ali H. 2016. Roles of MAS-related G protein coupled receptor-X2 (MRGPRX2) on mast cell-mediated host defense, pseudoallergic drug reactions and chronic inflammatory diseases. J Allergy Clin Immunol. 138(3):700–710. 10.1016/j.jaci.2016.04.051

Taguchi N et al. 2007. Mitotic Phosphorylation of Dynamin-related GTPase Drp1 Participates in Mitochondrial Fission. Journal of Biological Chemistry. 282(15):11521–11529. 10.1074/jbc.M607279200

Tatemoto K et al. 2006. Immunoglobulin E-independent activation of mast cell is mediated by Mrg receptors. Biochemical and Biophysical Research Communications. 349(4):1322–1328. 10.1016/j.bbrc.2006.08.177

Thrasher SM, Scalfone LK, Holowka D, Appleton JA. 2013. In vitro modeling of rat mucosal mast cell function in Trichinella spiralis infection. Parasite Immunol. 35(1):21–31. 10.1111/pim.12014

Trebak M, Kinet J-P. 2019. Calcium signalling in T cells. Nat Rev Immunol. 19(3):154–169. 10.1038/s41577-018-0110-7

Van Leeuwen MPC et al. 2015. Long-term efficacy of a 0.07% cetylpyridinium chloride mouth rinse in relation to plaque and gingivitis: a 6-month randomized, vehicle-controlled clinical trial. Int J Dent Hyg. 13(2):93–103. 10.1111/idh.12082

Varade D et al. 2005. Effect of salt on the micelles of cetyl pyridinium chloride. Colloids and Surfaces A: Physicochemical and Engineering Aspects. 259(1):95–101. 10.1016/j.colsurfa.2005.02.018

Voutchkova AM, Ferris LA, Zimmerman JB, Anastas PT. 2010. Toward molecular design for hazard reduction—fundamental relationships between chemical properties and toxicity. Tetrahedron. 66(5):1031–1039 (Advances in Green Chemistry). 10.1016/j.tet.2009.11.002

Wang Y et al. 2022. Dynamin-related protein 1 differentially regulates FcεRI- and substance P-induced mast cell activation. Journal of Allergy and Clinical Immunology. 150(5):1228–1231.e5. 10.1016/j.jaci.2022.04.028

Weatherly LM et al. 2016. Antimicrobial agent triclosan is a proton ionophore uncoupler of mitochondria in living rat and human mast cells and in primary human keratinocytes. J Appl Toxicol. 36(6):777–789. 10.1002/jat.3209

Weatherly LM et al. 2018. Antimicrobial Agent Triclosan Disrupts Mitochondrial Structure, Revealed by Super-resolution Microscopy, and Inhibits Mast Cell Signaling via Calcium Modulation. Toxicol Appl Pharmacol. 349:39–54. 10.1016/j.taap.2018.04.005

Weller SR et al. 2024. Antimicrobial cetylpyridinium chloride causes functional inhibition of mitochondria as potently as canonical mitotoxicants, nanostructural disruption of mitochondria, and mitochondrial Ca2+ efflux in living rodent and primary human cells. Food and Chemical Toxicology. 186:114547. 10.1016/j.fct.2024.114547

Wiemerslage L, Lee D. 2016. Quantification of Mitochondrial Morphology in Neurites of Dopaminergic Neurons using Multiple Parameters. J Neurosci Methods. 262:56–65. 10.1016/j.jneumeth.2016.01.008

Wong PT et al. 2014. Formulation and Characterization of Nanoemulsion Intranasal Adjuvants: Effects of Surfactant Composition on Mucoadhesion and Immunogenicity. Mol Pharmaceutics. 11(2):531–544. 10.1021/mp4005029

Zaitsu M et al. 2007. Estradiol activates mast cells via a non-genomic estrogen receptor-α and calcium influx. Mol Immunol. 44(8):1977–1985. 10.1016/j.molimm.2006.09.030

Zhang B et al. 2011. Human mast cell degranulation and preformed TNF secretion require mitochondrial translocation to exocytosis sites: Relevance to atopic dermatitis. Journal of Allergy and Clinical Immunology. 127(6):1522–1531.e8. 10.1016/j.jaci.2011.02.005

Zhang B et al. 2012. Mitochondria Distinguish Granule-Stored from de novo Synthesized Tumor Necrosis Factor Secretion in Human Mast Cells. International Archives of Allergy and Immunology. 159(1):23–32. 10.1159/000335178

Zhang Q et al. 2008. Hormesis and Adaptive Cellular Control Systems. Dose-Response. 6(2):dose-response. 10.2203/dose-response.07-028.Zhang

Zheng G, Schreder E, Sathyanarayana S, Salamova A. 2022. The first detection of quaternary ammonium compounds in breast milk: Implications for early-life exposure. J Expo Sci Environ Epidemiol. 32(5):682–688. 10.1038/s41370-022-00439-4

Zorova LD et al. 2018. Mitochondrial membrane potential. Anal Biochem. 552:50–59. 10.1016/j.ab.2017.07.009

