## Supplementary material for "Pharmaceutical agent cetylpyridinium chloride modulates mitochondrial dynamics to enhance calcium-stimulated immune mast cell function"

**Table of Contents**

**Fig. S1. Relative degranulation response of RBL mast cells exposed to various non-cytotoxic doses of Ca^2+^ ionophore A23187 for 1 hr..…...…………...…………….2**

**Fig. S2. Lactate dehydrogenase release cytotoxicity assay of combination of 300 nM A23187 dose and varying concentrations of CPC..…………………………………3**

**Fig. S3. Relative degranulation response of RBL mast cells exposed to various doses of CPC in conjunction with ionophore in galactose Tyrodes………….……...4**

**Fig. S4. Effects of CPC on ionophore-stimulated mast cell degranulation in galactose DMEM at varying levels of degranulation..…………………………………..5**

**Fig. S5. Linearity test for protein loading for Drp1 western blotting.………………..6**

**Fig. S6. CPC effects on Drp1 (Ser637) phosphorylation in unstimulated RBL mast cells assessed via western blot.……….…………………………………………………...7**

**Fig. S7. Fluorescence of MitoTracker Red FM treated with varying concentrations of CPC without cells…………………………………………………………………………..8**

**Fig. S8. Visual representation of steps for image analysis of mitochondrial translocation…………………………………………………………………………………....9**

**Fig. S9. Calcium ionophore and CPC effects on electron flow via Complex I…….10**

**Figure S1. Relative degranulation response of RBL mast cells exposed to various non-cytotoxic doses of Ca^2+^ ionophore A23187 for 1 hr.**

**Method:** RBL-2H3 mast cell degranulation was measured using the assay described in Materials & Methods, with the difference that various doses of Ca^2+^ ionophore were utilized. All treatment times were 1 hour. Assay was conducted in glucose-containing saline buffer BSA-Tyrodes.

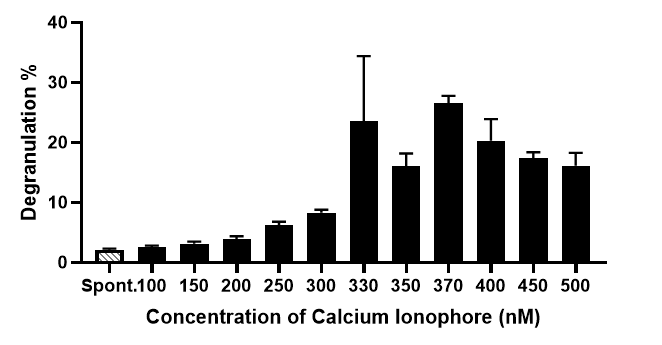

**Figure S1.** Spontaneous release cells ("Spont.") were exposed to BT in the absence of Ca^2+^ ionophore. Sample wells were exposed to 0 to 500 nM Ca^2+^ ionophore for 1 hr. Values shown are means displayed ± SEM, and represent at six independent days of experiments (each sample in triplicate).

**Result:** Calcium ionophore A23187 produced an increasing level of degranulation, starting with some stimulation over spontaneous release by about 150 nM and plateauing in the mid-300s range. At 300 nM, this lot of ionophore produced a degranulation response of ~7% in RBL-2H3 mast cells.

**Conclusion:** For the experiments portrayed in the manuscript’s figures, we utilized 300 nM Ca^2+^ ionophore. This dose balanced the need for sufficiently robust signaling over spontaneous levels with the goal of avoiding maximal or potentially cytotoxic doses. Also, by focusing our analyses on low levels of mast cell stimulation, we model physiologically relevant cell status (as opposed to a maximal anaphylactic state).
 **Figure S2. Lactate dehydrogenase release cytotoxicity assay of combination of 300 nM A23187 dose and varying concentrations of CPC.**

**Method:** Lactate dehydrogenase (LDH) activity release was measured in the treatment medium of RBL-2H3 cells in the same conditions as in degranulation (Fig. 2) and translocation/fission experiments (Figs. 4-8) (exposed to 300 nM Ca^2+^ ionophore & CPC), as described (Hutchinson et al. 2011).

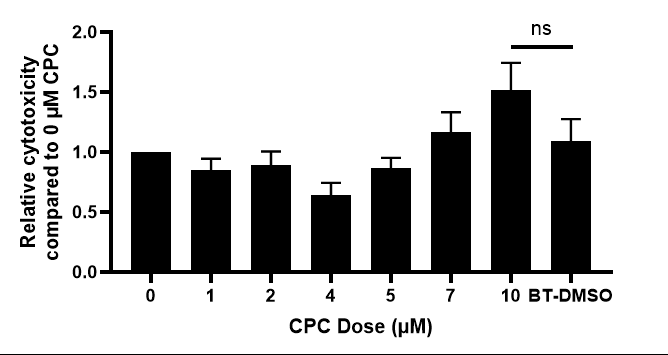

**Figure S2.** RBL-2H3 cells were treated with 300 nM Ca^2+^ ionophore along with 0 - 10 µM CPC to replicate conditions in degranulation experiments, in BT buffer. LDH release was measured using lactate dehydrogenase cytotoxicity assay (Cayman Chemical). The percentage cytotoxicity for each sample was normalized to the 0 µM CPC/300 nM ionophore control. “BT-DMSO” samples received neither ionophore nor CPC but did include the DMSO (0.0032%) vehicle. Values shown are mean ± SEM and represent data from eight independent days of experiments (triplicates per sample). Statistical significance was determined through one-way ANOVA with Tukey's post-hoc testing: ns = not significant.

**Result:** LDH cytotoxicity testing demonstrated no significant cell toxicity to the exposure conditions (Ca^2+^ ionophore, its vehicle, CPC, timing) utilized in this manuscript. Of note, all CPC/ionophore combinations used in experiments post-Fig. 2 in the manuscript use 2.5 µM CPC. Comparison of the BT-DMSO control group to the 0 µM CPC group shows that the 300 nM ionophore is not cytotoxic on its own. The combined treatments of 300 nM Ca^2+^ ionophore plus CPC were also not cytotoxic.

**Conclusion:** Any experimental effects caused by Ca^2+^ ionophore or Ca^2+^ ionophore + CPC in mast cells cannot be attributed to cytotoxicity.

**Figure S3. Relative degranulation response of RBL mast cells exposed to various doses of CPC in conjunction with ionophore in galactose Tyrodes.**

**Method:** RBL-2H3 mast cell degranulation was measured using the assay described in Materials & Methods, with the difference that galactose was used in place of glucose (still at 5 mM). Cells were kept in a glucose-containing complete media until the time of acute assay, during which hour they were supplied no glucose or other ready electron source. All treatment times were 1 hour. Assay was conducted in galactose-containing saline buffer BSA-Tyrodes.

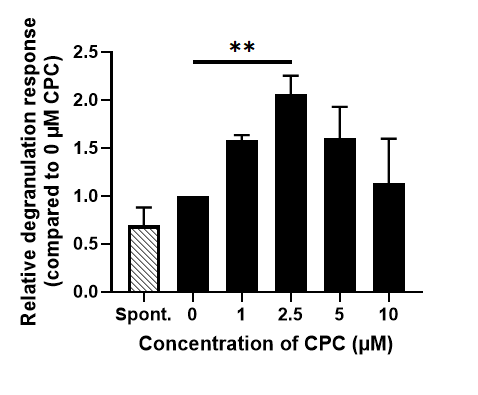

**Figure S3.** Spontaneous release cells ("Spont.") were exposed to BT in the absence of Ca^2+^ ionophore or CPC. Sample wells were exposed to 0 - 10 µM CPC along with 150-300 nM Ca^2+^ ionophore for 1 hr. Data are mean ± SEM and represent four independent experiments (samples in triplicate). Statistical significance was determined through one-way ANOVA with Tukey's post-hoc testing: ***p* < 0.05.

**Result:** CPC (1-5 µM) enhanced RBL-2H3 mast cell degranulation stimulated by Ca^2+^ ionophore. Low levels of ionophore (150 - 300 nM) were utilized in order to produce modest stimulation of degranulation (levels of 2.5 - 5.1%). The average ionophore-alone degranulation percentage was 3.4%, whereas the average spontaneous degranulation value was 2.2%. This condition (low stimulation of mast cell function) was needed, for causing CPC-induced enhancement in these low-energy conditions (galactose tends to be shunted to biosynthesis; no other electron sources are present in the buffer). At 2.5 µM, CPC maximally caused a 2-fold increase in degranulation over the control (compared to a boost of nearly 4-fold in Fig. 2, which is in the presence of glucose).

**Conclusion:** Galactose catabolism supplies the pentose phosphate pathway which creates biosynthetic precursors for molecules (fatty acids, cholesterol, DNA, and antioxidants among others) that support normal cellular function (Reitzer et al. 1979; TeSlaa et al. 2023). However, with no amino acids or other electron sources present in the buffer, the cells lack means to support the massive energy demands of degranulation. We found that, under these conditions of very low levels of stimulated degranulation in galactose-BT, CPC *still* stimulates ionophore-mediated degranulation.

**Figure S4. Effects of CPC on ionophore-stimulated mast cell degranulation in galactose DMEM at varying levels of degranulation.**

**Method:**

RBL-2H3 mast cell degranulation was measured using the assay described in Materials & Methods, with the difference that was conducted in glucose-free, galactose-containing DMEM media, as described in (Weatherly et al. 2013). All treatment times were 1 hour. Assay was conducted in galactose-containing saline buffer BSA-Tyrodes.

Immune cells may satisfy their energy needs with glycolysis (Crabtree 1928; Wang et al. 1976), thus bypassing mitochondrial oxidative phosphorylation; these are the conditions simulated in Fig. 2 of the main text. We also conducted degranulation experiments in DMEM containing both galactose and L-glutamine (but free of glucose), to force cells to utilize their mitochondria for energy production (Marroquin et al. 2007), in order to further explore the roles of energy status . In these conditions, L-glutamine is the primary source of cellular energy, which is catabolized to feed into the TCA cycle and subsequent oxidative phosphorylation, rendering glycolysis unnecessary. Experiments with galactose DMEM conditions probe whether CPC, in conjunction with ionophore stimulation, affects mast cell degranulation similarly to experiments conducted in BSA-Tyrodes (containing glucose). In these experiments, a range of Ca^2+^ ionophore doses (200 – 500 nM) was utilized to elicit a range of degranulation intensities (3.5% - 41.2% total degranulation) in RBL-2H3 mast cells.

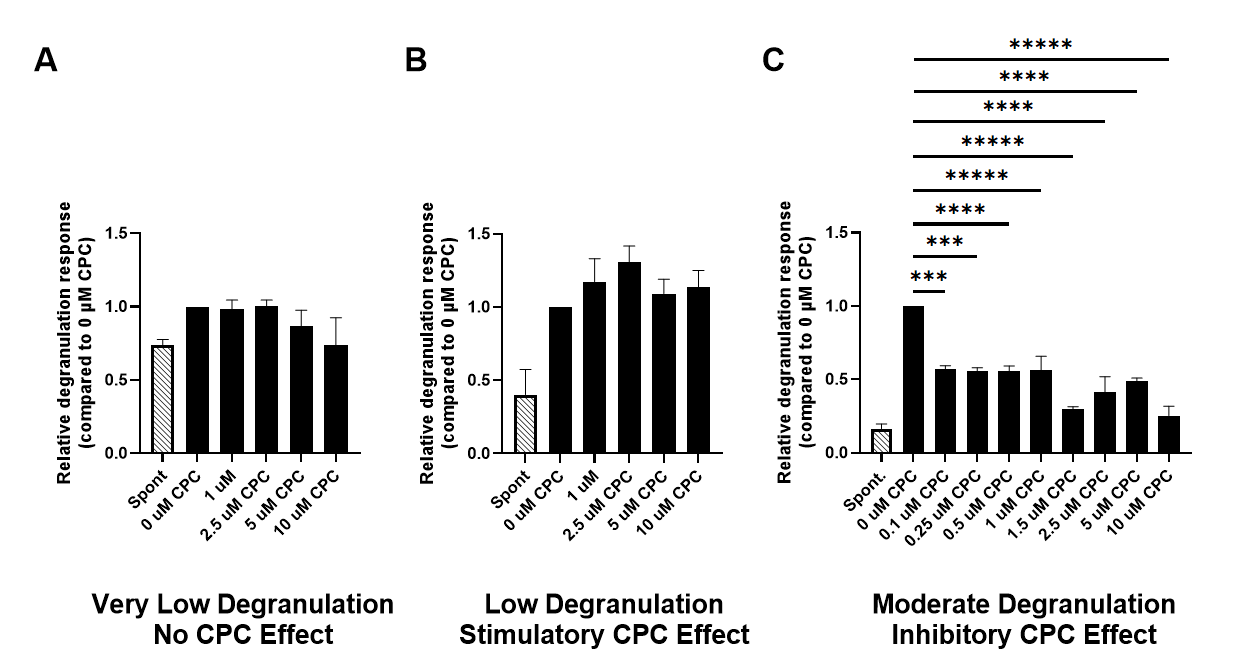

**Figure S4.** Spontaneous release cells ("Spont.") were exposed to galactose DMEM in the absence of Ca^2+^ ionophore or CPC. Sample wells were exposed to 0 - 10 µM CPC along with Ca^2+^ ionophore (range of doses noted in “Method”), for 1 hr. The assays were grouped based on very low (average 3.5%), low (average 10.8%), and moderate (average 20.4%) degranulation responses for the control (0 µM CPC with 200 - 500 nM ionophore). Values are mean ± SEM and represent at least two independent experiments for each group (each sample in triplicate). Statistical significance was determined through one-way ANOVA with Tukey's post-hoc testing: ****p* < 0.01, *****p* < 0.001, ******p* < 0.0001.

**Result:**

CPC had no effect on mast cells which were stimulated by ionophore but produced very low levels of granule exocytosis (Fig. S4A), on average 3.5%, a level only slightly above spontaneous degranulation in these experiments.

When cells were stimulated with ionophore and produced higher but still modest levels of degranulation (on average 10.8%) (Fig. S4B), CPC appears to have a stimulatory effect on degranulation with 2.5 µM CPC causing a 31% increase in degranulation over the control, although this increase did not reach statistical significance. Note that the CPC-induced pattern appears similar to that in Fig. 2 of the main text and also Fig. S3.

In cells stimulated with ionophore producing higher degranulation levels, averaging 20.4% (Fig. S4C), CPC drastically inhibited mast cell degranulation: just 0.1 µM CPC caused mast cell degranulation to decrease by 43% in cells with a moderate degree of degranulation.

**Conclusion:**

CPC effects on mast cell degranulation in glucose-free, galactose-containing DMEM are dependent on the extent of mast cell degranulation. with CPC causing no effect at very low levels of degranulation, some stimulation with low levels of degranulation, and inhibition at moderate levels of degranulation.

The lack of CPC effect in Fig. S4A is concurrent with our previously published finding that RBL cell spontaneous degranulation is unaffected by CPC (Obeng et al. 2023). This effect may be because at very low levels of degranulation, there may not be enough signaling for CPC to synergistically increase mast cell degranulation.

The reason for inability to reach significant stimulation in Fig. S4B may be related to the fact that, at the tested doses and in galactose DMEM media, CPC is known to inhibit mitochondrial ATP production (Weller et al. 2024) while it does not drastically inhibit ATP in the presence of glucose (Weller et al. 2024), the condition used in Fig. 2. Thus, while mitochondria undergo fission under these conditions (Weller et al. 2024), ATP is simultaneously inhibited, so cells do not have sufficient energy needed to harness the fissioned mitochondria for degranulation support. There is, still, an echo of the stimulation seen in Fig. 2 which is in glucose buffer, likely propelled by residual cellular energy.

Interestingly, in cells undergoing higher levels of degranulation (20.4% on average) in the galactose DMEM, very low doses of CPC actually inhibited mast cell degranulation. This severe inhibition may be explained by the known, strong inhibition of mitochondrial ATP production due to CPC when in galactose DMEM (Weller et al. 2024). CPC is very mitotoxic; for example, CPC inhibits electron flow into Complex IV of the electron transport chain with similar potency as cyanide (Ledue et al. 2026). Therefore, in the galactose DMEM condition in which the only pathway cells have to create energy (oxidative phosphorylation) is potently inhibited by CPC, mast cells are unable to meet the huge energy demands needed for degranulation when higher levels of ionophore-induced cell stimulation happen. This potent mitochondrial ATP inhibition then outweighs the stimulatory effect of CPC-induced fission-supported mitotranslocation.

**Figure S5. Linearity test for protein loading for Drp1 western blotting.**

**Method:**

A linearity test was done to determine the range within which Drp1 signal increases linearly with increasing amounts of loaded protein, in order to determine the appropriate amount of protein to load in each lane of the gel. Published methods (Obeng et al. 2024) were employed. Cell preparation, protein concentration determination of cell lysates, and western blotting were performed as in “Materials and Methods,” except that variable quantities of protein sample were assessed. After imaging, a curve was generated with “mean fluorescence intensity,” as represented by integrated density in Fiji ImageJ, on the y-axis and amount of protein on the x-axis for the target of interest.

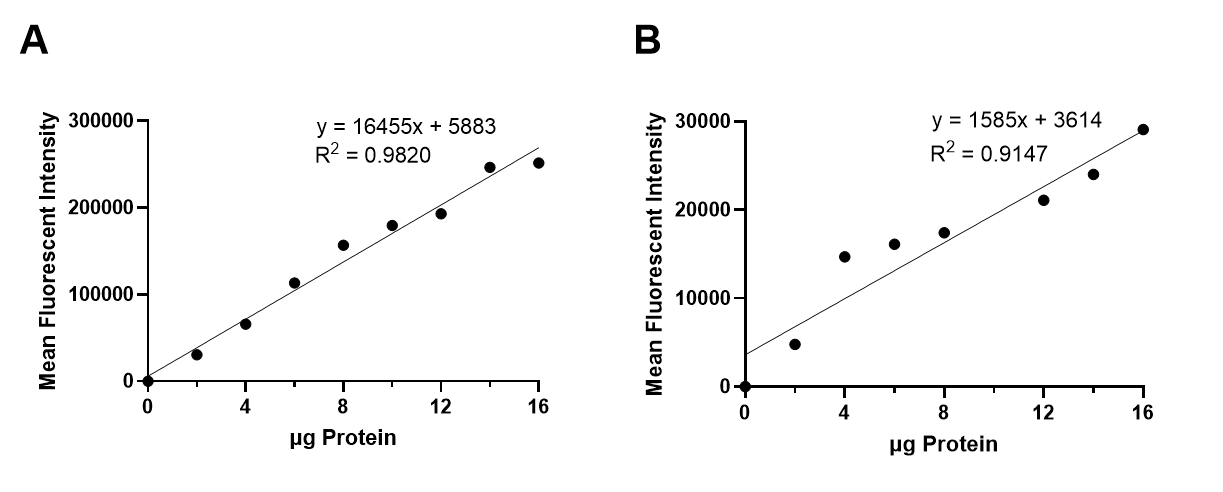

**Figure S5.** Linearity test for total Drp1 antibody **(A)** and phospho-Drp1 (Ser616) **(B).** Varying amounts of RBL-2H3 mast cell whole cell lysates were probed with total Drp1 or phospho-Drp1 (Ser616) antibodies and a corresponding fluorescent antibody.

**Result:** Linearity is represented by R^2^ value, which is 0.982 for (**A**) and 0.9147 for (**B**).

**Conclusion:** Calculated R^2^ values of 0.982 (A; total Drp1) and 0.9147 (B; phospho-Drp1 (Ser616)) indicate good linearity of the antibody for the µg of protein used. All western blot experiments examining phosphorylation levels of Drp1 (Ser616) utilized protein amounts within this linear range (8 - 12 µg).

**Figure S6. CPC effects on Drp1 (Ser637) phosphorylation in unstimulated RBL mast cells assessed via western blot**

**Method:** Cells were treated and proteins were probed using the western blot protocol described in Materials & Methods for the Drp1 Ser616 experiments, with the following modifications: phospho-Drp1 Ser637 (Cell Signaling Technologies) was used. Cells were treated with 0, 2.5 µM, or 5 µM of CPC for 1 hour. Protein bands (or equivalent-size boxes from the total Drp1 side of the immunoblot) of Ser637-phosphorylated Drp1 were normalized to total Drp1 in each sample (control of 0 µM CPC, 2.5 µM CPC, and 5 µM CPC).

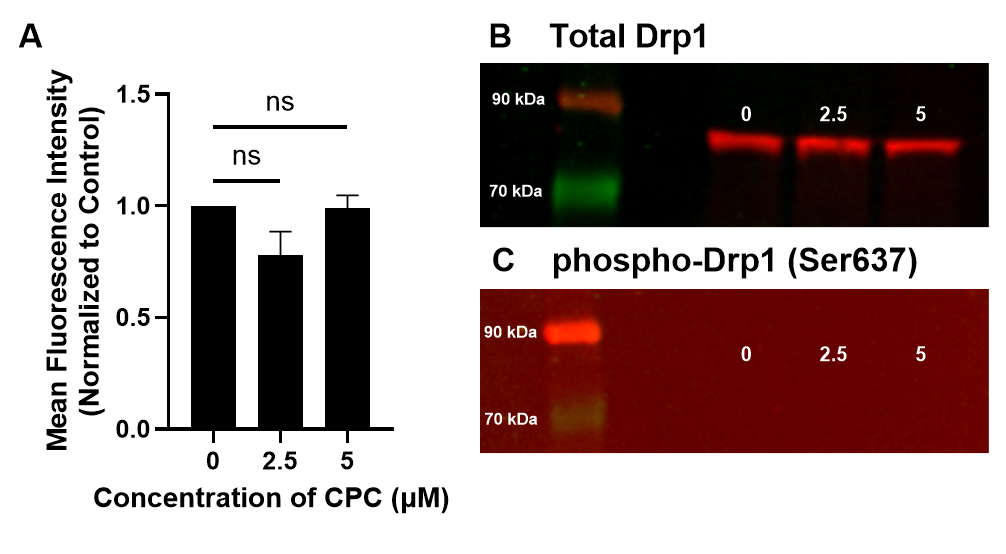

**Figure S6.** Cells were treated with 0 µM, 2.5 µM, or 5 µM of CPC for 1 hr before lysis, then electrophoresed, immunoblotted, and probed with total Drp1 or phospho-Drp1 (Ser637) antibodies. On each day, the phospho-Drp1 signal was normalized to the corresponding total Drp1 signal. **(A)** Data are mean ± SEM and represent three independent experiments normalized to the control group (0 µM). Statistical significance was determined through one-way ANOVA via Turkey’s post-hoc testing: ns = not significant. **(B)** and **(C)** are representative images of experiments. Ladder shows molecular weight in kDa.

**Result:** In some experiments, no band was detected at the appropriate molecular weight in the Drp1 (Ser637) phospho-blot; in others, a faint band was present but did not change with CPC exposure. Thus, phosphorylation of Drp1 at Ser637 seems low, even undetectable, in RBL-2H3 mast cells, even following CPC exposure.

**Conclusion:** Phosphorylation at Drp1 Ser637 is very low or undetectable in RBL-2H3 mast cells that are resting or treated with CPC. We found that CPC does not affect phosphorylation of Drp1 Serine 637 at the doses tested (2.5 µM, and 5 µM CPC, each for 1 hr).

**Figure S7. Fluorescence of MitoTracker Red FM treated with varying concentrations of CPC without cells.**

**Method:** Various doses (0 µM, 2 µM, 5 µM, 10 µM, and 20 µM) of CPC were made in BT. Also in BT, Mitotracker Red FM (ThermoFisher) was prepared using DMSO vehicle, for a final experimental concentration of 10 nM and 0.0025% DMSO. The 10 nM dose of Mitotracker Red FM was used to estimate the concentration in cellular experiments, following application and washing. Samples were placed into a 96-well black bottom plate (Greiner CellStar). BT with 0.0025% DMSO was added to the background wells to ensure DMSO levels were the same across all wells. The doses of CPC were added to the treatment wells for final concentrations of 0 µM, 1 µM, 2.5 µM, 5 µM, and 10 µM. Background 1 received a final concentration of 0 µM CPC in BT with 0.0025% DMSO, while Background 2 received a final concentration of 10 µM CPC with 0.0025% DMSO. The plate was then covered and left to incubate at 37°C for 1 h. At 1 h, the fluorescence was read on a microplate reader (Synergy 2, BioTek), excitation 530 nm, emission 645 nm.

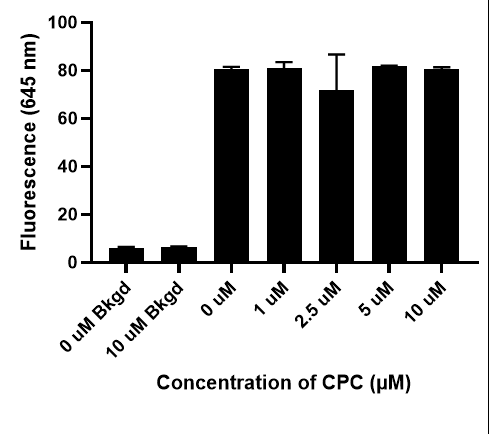

**Figure S7.** Fluorescence was measured after 10 nM MitoTracker Red FM in BSA-Tyrodes was exposed to CPC for 1 h. Background wells contained no MitoTracker Red FM and either 0 µM CPC or 10 µM CPC.

**Result:** The tested 10 nM Mitotracker Red FM dye produced a robust signal in the plate reader, despite the absence of cells; this signal was higher than background. CPC had no effect on background signal at the excitation and emission wavelengths needed to assess this probe. No dose of tested CPC significantly altered the fluorescence of Mitotracker Red FM, in comparison to the control.

**Conclusion:** CPC does not interfere with the fluorescent emission of Mitotracker Red FM. Thus, observed experimental effects of CPC on Mitotracker Red FM fluorescence are likely physiological in nature.

**Figure S8. Visual representation of steps for image analysis of mitochondrial translocation.**

**Method:** For mitochondrial translocation analysis, first, a cell is selected and magnified for analysis such that only one cell is in the field of view.  In ionophore- or antigen-treated samples (though not “Spont.” controls), only degranulating cells were visually selected for analysis (via presence of morphological features such as actin ruffling or spreading), in order to focus analysis on cells undergoing active signaling.  Next, a duplicate hyperstack is created of that cell including the MitoTracker Red FM channel and the nuclear Hoechst channel. Two masks are created, of both the mitochondria and nuclear channels. The nuclear channel is dilated by 30 pixels, and holes in the mask are filled in. Any bubbles due to pixels outside of the nucleus being dilated are removed. This dilated, 8-bit nuclear mask is used to create a selection and measure the mitochondrial area (number of pixels with MitoTracker Red FM) inside this perinuclear space. Mitochondrial area was measured via the raw integrated density of this mitochondrial mask for each cell, both within the perinuclear region and, separately, in the whole cell (whole field of view). Raw integrated density is a proxy for area covered by the mitochondria (though not number of mitochondria per pixel) because each pixel is either binary white (255) indicating mitochondrial presence or black (0) indicating no mitochondrial presence. Finally, the perinuclear mitochondrial area was divided by the whole cell mitochondrial area, subtracted from 1, and multiplied by 100 to gain the percentage of mitochondria localized outside of the perinuclear region (% mitochondrial translocation). The code for this macro is included below. The “Read and Write Excel” plugin is needed to record results to an Excel file.
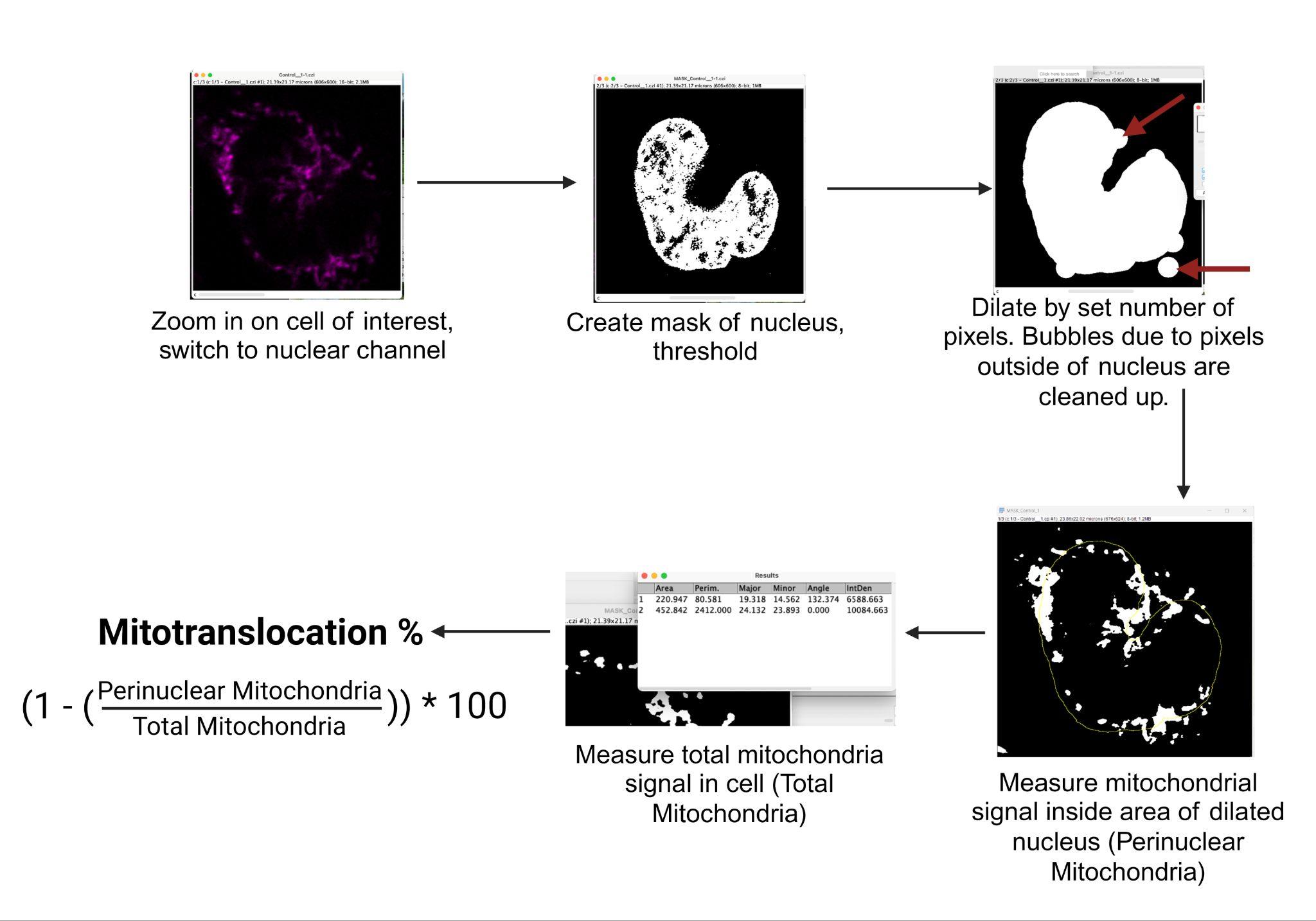

**Figure S8.** A visual representation of the macro used to measure mitochondrial translocation, created in Fiji ImageJ2.

**Mitotranslocation Macro Code:**

// Select a cell and duplicate the hyperstack of selected cell

waitForUser("Make a rectangular selection around a cell in the Nucleus (Hoechst) channel...");

run("Duplicate...", "title=dupe duplicate");

// Thresholds nucleus & creates mask

setAutoThreshold("Default dark no-reset");

run("Threshold...");

setOption("BlackBackground", true);

run("Convert to Mask", "background=Dark calculate black create");

// Fills holes and enlarges nucleus

run("Fill Holes", "slice");

run("Maximum...", "radius=30 slice");

waitForUser("Use the pencil tool (set to Black) to erase any particles which are not part of the primary nucleus...");

// Creates selection around enlarged nucleus and measures the selection in the mitochondrial channel, then measures all mitochondria in the cell

// Records results in an Excel spreadsheet

run("Create Selection");

Stack.setChannel(1);

run("Measure");

run("Read and Write Excel", "file=[/File_path/file_name.xlsx] sheet=Perinuc_Mito_Results stack_results dataset_label=Perinuc_Mito_Results");

run("Clear Results");

run("Select None");

run("Measure");

run("Read and Write Excel", "file=[/file_path/file_name.xlsx] sheet=All_Mito_Results stack_results dataset_label=All_Mito_Results");

run("Clear Results");

// Clears results

// Closes any windows not needed after measurement of cell

close("Results");

close("dupe");

close("MASK_dupe");

close("Threshold");

**Figure S9. Calcium ionophore and CPC effects on electron flow via Complex I**

**Method:** Electron flow from NADH through the mitochondrial electron transport chain was measured in RBL-2H3 cells using an NADH-Glo kit (Promega) as published (Ledue et al. 2026) with the difference that cells were plated in phenol red-free RBL-2H3 media, and the 1 hr exposures to CPC and ionophore were performed in BSA-Tyrodes buffer. A higher ionophore dose was used, compared to other portions of this manuscript, due to issues of potency.

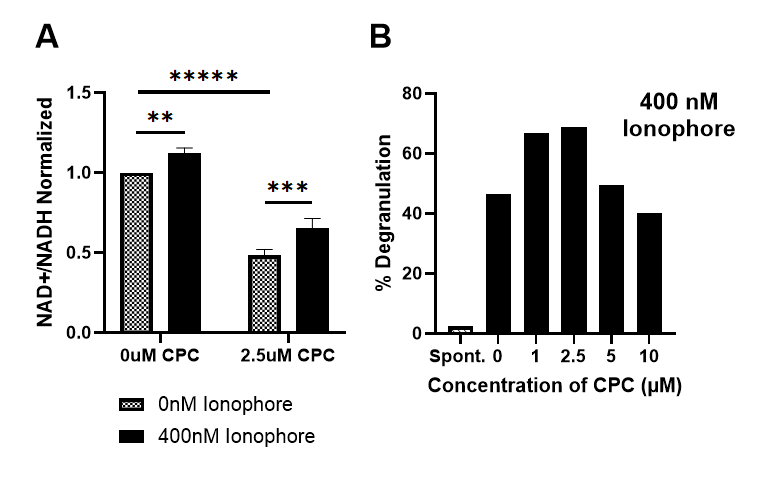

**Figure S9. (A)** A luminescence-based NADH-Glo assay was used to measure the NAD+/NADH ratio in intact RBL-2H3 cells in glucose-containing BT. Ratios (NAD+/NADH) were normalized to the trial lacking CPC and ionophore (0 µM and 0 nM, respectively). Data are mean ± SEM and represent four independent experiments, each with three technical replicates. Significance was determined with a two-way ANOVA and uncorrected Fisher's LSD multiple comparisons test in GraphPad Prism. ** p < 0.05, ***p < 0.01, *****p < 0.0001. **(B)** represent the % total degranulation of mast cells exposed to the same dose of Ca^2+^ ionophore, during the same exposure conditions and duration under which the NADH-Glo experiments were conducted. Data in the degranulation graph represents one independent experiment.

**Results:**

The utilized dose of ionophore (400 nM) produced a similar pattern of CPC stimulation of degranulation, as reported with 300 nM (Fig. 2). The potential effects of CPC on the luminescent probe were tested: it was determined that CPC does not interact directly with the probe (data not shown).

Between the 0 nM ionophore trials, CPC inhibited NADH oxidation, a measurement associated with complex I activity, by half. This inhibition is consistent with what is seen in (Ledue et al. 2026), where concentrations of CPC as low as 250 nM cut electron flow through Complex I by roughly half.

For trials with either 0 or 2.5 µM CPC, Ca^2+^ ionophore caused a small but significant stimulation in NADH oxidation. For trials with 2.5 µM CPC, stimulation of NADH oxidation caused by Ca^2+^ ionophore was noted, although it was not enough to fully recover from the CPC effect.

**Conclusion:** Calcium ionophore dosing leads to stimulation of electron flow through Complex I, even as CPC is inhibitory. Increased flow of electrons through the ETC may be related to the stimulation of mitochondrial membrane potential (Fig. 8A) seen in CPC/Ionophore trials – but this NADH oxidation effect does not fully explain the Fig. 8A findings. While there are greater ionophore stimulation effects on NADH oxidation in the presence of CPC, it is insufficient for full recovery of NADH oxidation rates from the CPC inhibition. Thus, increased ETC activity, at least at Complex I, is likely not the sole cause of the MMP effect.

**References**

Crabtree HG. 1928. The carbohydrate metabolism of certain pathological overgrowths. Biochem J. 22(5):1289–1298. https://doi.org/10.1042/bj0221289

Hutchinson LM et al. 2011. Inorganic arsenite inhibits IgE receptor-mediated degranulation of mast cells. J Appl Toxicol. 31(3):231–241. https://doi.org/10.1002/jat.1585

Ledue EL et al. 2026. Antimicrobial cetylpyridinium chloride disrupts the mitochondrial electron transport chain as potently as cyanide, via cardiolipin interference at cytochrome C. 2026.08.18.745331 [accessed 2026 Nov 9]. https://www.biorxiv.org/content/10.64898/2026.08.18.745331v1. https://doi.org/10.64898/2026.08.18.745331

Marroquin LD et al. 2007. Circumventing the Crabtree Effect: Replacing Media Glucose with Galactose Increases Susceptibility of HepG2 Cells to Mitochondrial Toxicants. Toxicological Sciences. 97(2):539–547. https://doi.org/10.1093/toxsci/kfm052

Obeng B et al. 2023. Pharmaceutical agent cetylpyridinium chloride inhibits immune mast cell function by interfering with calcium mobilization. Food and Chemical Toxicology. 179:113980. https://doi.org/10.1016/j.fct.2023.113980

Obeng B et al. 2024. Anti-microbial cetylpyridinium chloride suppresses mast cell function by targeting tyrosine phosphorylation of Syk kinase. J Immunotoxicol. 21(1):2443397. https://doi.org/10.1080/1547691X.2024.2443397

Reitzer LJ, Wice BM, Kennell D. 1979. Evidence that glutamine, not sugar, is the major energy source for cultured HeLa cells. Journal of Biological Chemistry. 254(8):2669–2676. https://doi.org/10.1016/S0021-9258(17)30124-2

TeSlaa T, Ralser M, Fan J, Rabinowitz JD. 2023. The pentose phosphate pathway in health and disease. Nat Metab. 5(8):1275–1289. https://doi.org/10.1038/s42255-023-00863-2

Wang T, Marquardt C, Foker J. 1976. Aerobic glycolysis during lymphocyte proliferation. Nature. 261(5562):702–705. https://doi.org/10.1038/261702a0

Weatherly LM, Kennedy RH, Shim J, Gosse JA. 2013. A microplate assay to assess chemical effects on RBL-2H3 mast cell degranulation: effects of triclosan without use of an organic solvent. J Vis Exp. (81):e50671. https://doi.org/10.3791/50671

Weller SR et al. 2024. Antimicrobial cetylpyridinium chloride causes functional inhibition of mitochondria as potently as canonical mitotoxicants, nanostructural disruption of mitochondria, and mitochondrial Ca2+ efflux in living rodent and primary human cells. Food and Chemical Toxicology. 186:114547. https://doi.org/10.1016/j.fct.2024.114547
